# A core genetic mechanism integrates growth hormone signals to control meristem fate and inflorescence architecture in setaria and maize

**DOI:** 10.1101/2025.06.27.660957

**Authors:** Jaspreet Sandhu, Jiani Yang, Max Braud, Edoardo Bertolini, Andrea L Eveland

## Abstract

Inflorescence architecture in cereals is a major determinant of grain yield and harvestability. Architecture is determined by the number, arrangement and order of lateral branches, which are formed from clusters of stem cells called meristems. Variation in branching patterns arises from regulation of meristem determinacy, i.e., indeterminate meristems maintain meristematic activity allowing for higher order branching and determinate meristems form a terminal organ. The timing by which an indeterminate branch meristem acquires a determinate fate and transitions to a spikelet meristem (SM), ultimately forming the grain-bearing spikelet, defines architecture and grain yield potential. This study leverages the unique inflorescence morphology of model grass *Setaria viridis* (setaria) to identify factors controlling SM fate and determinacy and characterize their interactions. In setaria, axillary branches terminate either in a spikelet or sterile bristle, where the latter loses SM identity due to brassinosteroid (BR)-mediated disruption of organ boundary formation. A forward genetics screen identified two mutants with only bristles on their panicles; one defective in gibberellic acid (GA) signaling caused extreme elongation and disruption of spikelet development and the other homeotically converted spikelets to bristles. The latter named *spikeletless* (*spkl*) was mapped and validated as an ortholog of the maize determinacy factor *ramosa1 (ra1)*. Genetic analyses showed that *SvRa1* acts upstream of BR-mediated regulation of SM identity and determinacy. A model is proposed where SvRA1 maintains SM fate by modulating GA homeostasis, spatially restricting BR signaling and preserving boundary gene expression and determinacy. In bristle primordia, which are generally paired with spikelets, GA and BR act synergistically. Comparative analyses in maize highlight pathway conservation in inflorescence architecture with some specific variations.

## Introduction

Cereal crops provide more than 50% of humanity’s caloric intake and therefore have an enormous impact on food security worldwide. With a world population expected to reach 9.7 billion by 2050, accelerated development of high-yielding crop varieties is needed to meet future food supply demands. In cereals, grains are borne on inflorescences, which display complex branching patterns that vary across the grass family, Poaceae. These diverse inflorescence architectures are determined by the placement, number and orientation of lateral branches that are initiated from pools of pluripotent stem cells called meristems (Tanaka et al. 2013; Hong and Fletcher 2023). Inflorescence architecture is a major determinant of yield by influencing grain number and size, harvestability, and reproductive success. Therefore, knowledge of the mechanisms controlling meristem identity and determinacy during inflorescence development can be harnessed for enhancing yield potential (Sakuma and Schnurbusch 2020; Koppolu et al. 2022; Lindsay et al. 2024).

During development, each meristem has three possible fates: it can remain meristematic and produce lateral meristems on its flanks (indeterminate), it can be terminally differentiated into an organ such as a flower (determinate), or it can simply cease development (Bommert and Whipple 2018; Tanaka et al. 2013). In grass inflorescences, an indeterminate inflorescence meristem (IM) kicks off axillary meristems until itself becomes determinant or ceases development. These axillary meristems are typically indeterminate branch meristems (BMs), which ultimately transition to a determinant spikelet meristem (SM) that forms a spikelet. The spikelet is the terminal structure that marks cessation of a branching event and bears the flowers and ultimately grains. The timing of the transition from BM to SM, i.e., from indeterminate to determinate growth, underlies the extensive variation in branching patterns found across the grasses (Kellogg 2001, 2022; McSteen and Kellogg 2022). This shift from BM to SM involves the precise expression of spikelet identity genes and the establishment of organ boundaries, domains of reduced growth that spatially restrict organ identity and determinacy factors from meristem maintenance signals (Žádníková and Simon 2014; Richardson and Hake 2018). Several factors are involved in setting up boundaries, including boundary-specific transcription factors (TF)s and the exclusion of growth promoting hormones like brassinosteroids (BRs) and gibberellic acid (GA) (Gendron et al. 2012; Arnaud and Laufs 2013; Hepworth and Pautot 2015). BRs, for example, inhibit the expression of boundary promoting TFs e.g., *CUP-SHAPED COTYLEDON (CUC)* and *LATERAL ORGAN FUSION (LOF)* through the BR-responsive TF BRASSINAZOLE-RESISTANT 1 (BRZ1), thus establishing antagonistic signaling gradients that demarcates boundaries (Gendron et al. 2012; Bell et al. 2012). BR and GA both accumulate in the developing lateral primordia where low ratios of these hormones maintain determinate fate (Hepworth and Pautot 2015).

SM identity is marked by the initiation of lateral glume primordia (Wang et al. 2022) and expression of a conserved AP2 TF, initially characterized in *Zea mays* (maize) encoded by the *branched silkless 1 (bd1*) gene (Chuck et al. 2002). BD1 establishes SM identity and determinacy by suppressing the meristems in axils of subtending glumes. Orthologs of *bd1* have been characterized in several other grass species and are known by different names: *Oryza sativa* (rice) *FRIZZY PANICLE* (*FZP*) (Komatsu et al. 2003), *Brachypodium distachyon MORE SPIKELETS 1* (*MOS1) (Derbyshire and Byrne 2013)*, and *Hordeum vulgare* (barley) *COMPOSITUM 2* (*COM2*) (Poursarebani et al. 2015). In *Triticum aestivum* (wheat), mutations in *COM2* enhanced supernumerary spikelet production leading to increased grain yield (Dobrovolskaya et al. 2015) and underlies the “Miracle-Wheat” phenotype (Poursarebani et al. 2015). In rice, moderate suppression of FZP by BZR1 delayed determinate fate just enough to produce more spikelets and increased yield (Wang et al. 2020; Huang et al. 2018; Bai et al. 2017).

In Andropogoneae grasses including maize and sorghum, spikelets are borne in pairs and BMs transition to a spikelet pair meristem (SPM) prior to making two SMs (McSteen 2006; Whipple 2017). The C2H2 TF RAMOSA 1 (RA1) is required for SPM determinacy and repression of branching. In maize, expression of *ra1* is consistent with the abrupt shift from making long to short branches in the tassel, whereas in sorghum and *Miscanthus* that make numerous higher order branches along the inflorescence, *RA1* is induced later and at lower levels (Vollbrecht et al. 2005). The paired spikelet character appears to have arisen independently in the Paniceae, a sister clade to the Andropogoneae (McSteen 2006; Vollbrecht et al. 2005). Within the Paniceae, “bristle grasses” produce sterile bristles that are modified branches and appear to be paired with fertile spikelets (Doust and Kellogg 2002; Yang et al. 2018). Among these, *Setaria viridis* (green millet; referred to here as setaria) is a weedy grass and the wild relative of domesticated cereal *Setaria italica* (foxtail millet). Setaria has emerged as a useful model system for translational studies to closely related cereals such as maize and sorghum (Huang et al. 2016).

We previously reported that mutations in a gene encoding a rate limiting BR biosynthesis enzyme CYP450 homeotically converted bristles to spikelets resulting in a “bristleless” inflorescence, which was phenocopied by treatment with a BR inhibitor (Yang et al. 2018). Studies of this *bristleless 1* (*bsl1*) mutant revealed that BRs were required for the acquisition of bristle fate in one of two paired SMs, which initially both expressed SM identity genes (Yang et al. 2018). This homeotic switch in axillary meristem fate provides an ideal system for dissecting the molecular mechanisms controlling SM determinacy pathways in important cereal crops. Here, we leveraged this system in setaria to identify additional components in the bristle vs spikelet fate program in a forward genetics screen, tested genetic interactions among them, and evaluated transcriptional profiles underlying bristle vs spikelet development. We identified the ortholog of RA1 as a key regulator upstream in this fate decision and propose a model by which gradients of growth hormones maintain a dichotomous pattern of determinacy/indeterminacy in the spikelet-bristle pair. Finally, we evaluated comparable genetic interactions in maize.

## Results

### *SvSlr1* is the functional ortholog of rice *SLENDER1,* which encodes a negative regulator of GA signaling

M_2_ families from an N-Nitroso-N-methylurea (NMU) mutagenized population of *Setaria viridis* (Huang et al. 2017; Yang et al. 2018) were screened for mutants with defects in bristle vs spikelet fate. Two mutants were identified that produced few or no spikelets on the panicle, making only bristles (Fig. S1). One mutant we named *spikeletless* (*spkl*) made no spikelets except for a few at the tip and bristles were densely packed along the panicle (Fig. S1, Table S1). The second mutant produced very elongated panicles with only bristles that were also very elongated. The whole-plant phenotypes of this mutant bore striking resemblance to those of the rice *slender1* (*slr1*) mutant (Fig. 1A, (Ikeda et al. 2001)), including narrower but more elongated leaves and stems as well as the elongated panicles (Fig. 1B, Table S1). We therefore named this mutant *sv slender1* (*svslr1*). The *svslr1* mutants produced fewer main tillers than segregating normal siblings (Fig. 1A, Table S2) but developed multiple secondary tillers on the nodes of primary tillers (Fig. S2).

**Fig. 1.**
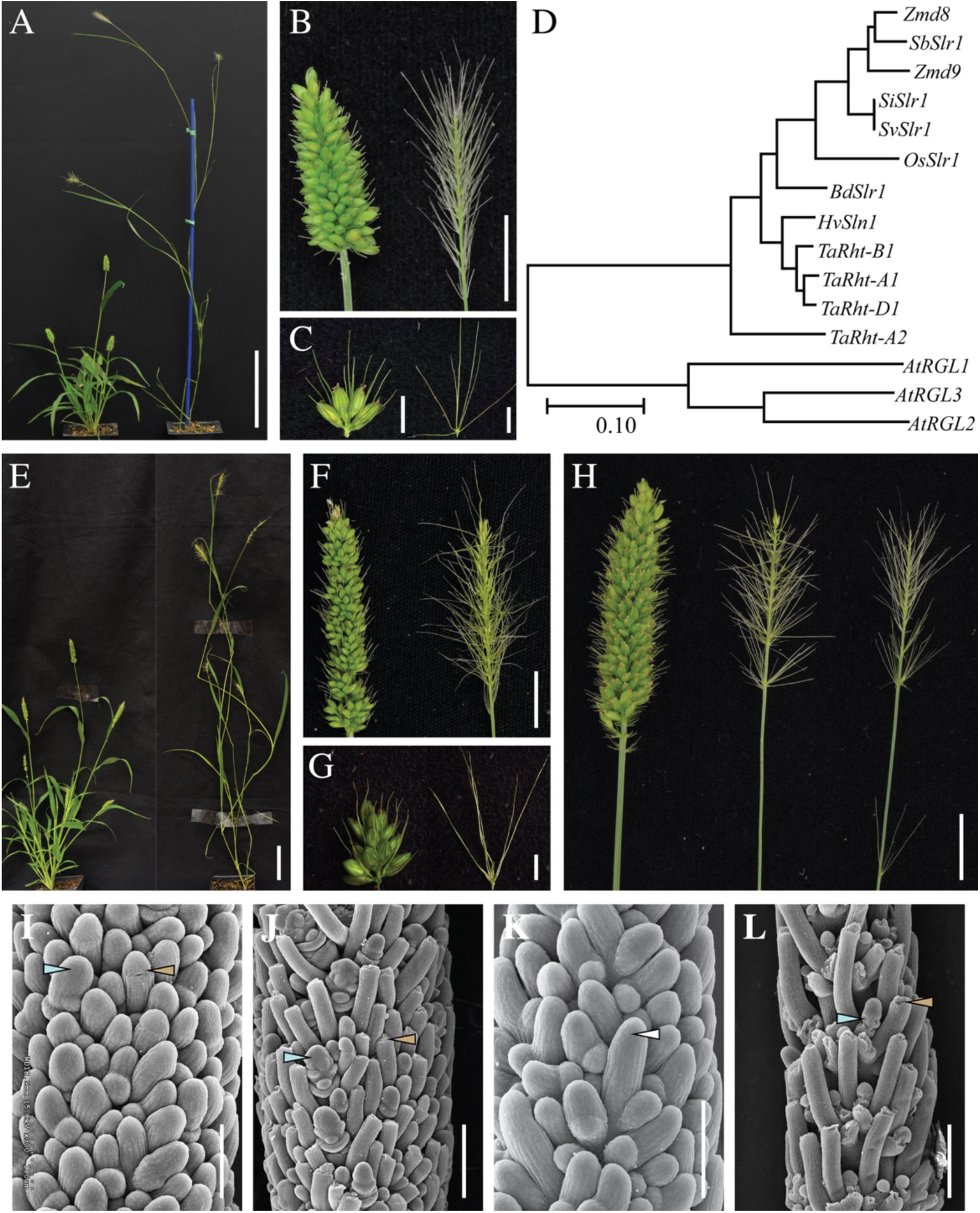
A mutation in *DELLA* results in hyper-elongation and panicles with all bristles. (A-C) Whole plant and panicle phenotypes of *svslr1* (right) compared to parental genotype, A10.1 (left). (A) Compared to A10.1, *svslr1* showed hyper-elongated stature, narrower leaves and (B) elongated panicles. (C) Primary branch clusters of A10.1 (left) and *svslr1* (right) showed lack of spikelets and elongation of bristles in *svslr1*. (D) A phylogenetic tree of *Slr1*-like coding sequences in selected grasses*. Setaria virdis* (*Sv*)*, Setaria italica* (*Si*)*, Sorghum bicolor* (*Sb*)*, Hordeum vulgare* (*Hv*) and *Brachypodium distachyon* (*Bd*) have single *Slr1* genes while *Zea mays* (*Zm*) and *Triticum aestivum* (*Ta*) have multiple copies. The coding sequences were used to construct a phylogenetic construct tree in MEGA X (Maximum Likelihood method and Tamura-Nei model). (E-G) CRISPR (CR)-cas9 based editing of *SvSlr1* validated the all-bristle phenotype of *svslr1*. (E) Plants, (F) panicles and (G) primary branch clusters of *cr-slr1* (right) compared to parental genotype, ME034 (left). (H) Panicles from untreated A10.1 (left), GA_3_-treated A10.1 (middle) and untreated *svslr1* (right) plants show that exogenous GA₃ treatment of A10.1 phenocopies the all-bristle phenotype of *svslr1*. (I-L) Scanning electron microscopy of developing (I-J) A10.1 and (K-L) *svslr1* panicles. (I) At 16 days after sowing (DAS), A10.1 branch meristems (BMs) were mostly undifferentiated, except for a few visible spikelet meristems (SMs, blue arrow). A few elongated meristems, likely precursors of bristles, with an indented ring (gold arrow) around the meristem tip were also observed. (J) Bristles with a severed meristem at the indented ring (gold arrow) were fully formed and SMs (blue arrow) had transitioned to floral meristems (FMs) around 18 DAS in A10.1. (K) In *svslr1,* elongated BMs (white arrow) with intact meristem were already visible at 14 DAS. (L) The *svslr1* panicles were more advanced than A10.1 as fully developed and elongated bristles (gold arrow) were already formed in *svslr1* around 15 DAS. SMs appear to abort (blue arrow) before differentiation in *svslr1*. **Scale bars:** 10 cm (A), 1 cm (B, F, H), 2 mm (C, G), 5 cm (E), 100μm (I, K) and 200 μm (J, L).

Compared to normal A10.1 siblings where spikelets and bristles are largely paired, primary branch clusters of *svslr1* mutant panicles produced only bristles, occasionally with aborted spikelets that failed to make seeds (Fig. 1B-C, S1). Density of primary branch clusters was decreased on s*vslr1* panicles compared to normal siblings (Table S1). Since homozygous *svslr1* does not produce seeds, the mutant stock was maintained as heterozygous. Progeny of selfed heterozygotes segregated in 3:1 normal: *svslr1* ratio, suggesting that *svslr1* is a single locus recessive allele (95:33; P (χ2, 1 d.f.) =0.8383).

The rice *slr1* phenotype results from a null mutation in the single copy *SLR1* gene that encodes DELLA, a negative regulator of GA signaling (Ikeda et al. 2001). A phylogenetic analysis of *SLR1-like* (*DELLA*) genes in grasses, using *Arabidopsis thaliana* as an outgroup, indicated that several grass species, including setaria, have a single copy of *DELLA* like in rice (Fig. 1D). We sequenced the single copy *SvSlr1* gene (*Sevir. 9G121800*) from several *svslr1* mutant individuals and compared the sequences to the A10.1 reference genome (Mamidi et al. 2020). A single nucleotide polymorphism (SNP) (C to A) 843 bp downstream of the ATG in *SvSlr1* changed a Tyrosine to a premature stop codon in *svslr1* mutants (Fig. S2). A Cleaved Amplified Polymorphic Sequences (CAPS) marker was used to genotype segregating populations - 18 and 63 individuals with *svslr* and wild-type phenotypes, respectively – and showed 100% linkage between genotype and *slender* phenotype (Fig. S2). To functionally validate this mutation, we used CRISPR/Cas9-based gene editing to generate *cr-slr1* mutants in the ME034 “transformable” setaria genotype (Thielen et al. 2020) where a 1 bp deletion 385 bp downstream of the ATG resulted in a frameshift mutation (Fig. S3). Edited *cr-slr1* plants phenocopied vegetative and panicle phenotypes of the original *svslr1* mutant (Fig. 1E-G).

In rice *slender* mutants, loss of function mutations in the GRAS domain cause a constitutive GA response due to lack of DELLA-mediated repression of GA signaling. We hypothesized that the all-bristle phenotype observed in *svslr1* was caused by hyper-active GA response. To test this, we treated setaria A10.1 plants with 10 mM GA_3_ at 14 DAS (just prior to spikelet and bristle differentiation) for a week.

GA_3_-treated A10.1 panicles phenocopied those of *svslr1* (Fig. 1H) as well as whole-plant vegetative phenotypes including hyper-elongation and lack of tillers (Fig. S4). According to published RNA-seq data from a setaria inflorescence developmental series (Zhu et al. 2018), expression of *SvSlr1* was lowest when terminal BMs poised to become a bristle or SM became morphologically distinct (15 DAS), and highest when SMs differentiated into FMs (17 DAS; Fig. S5).

### Spikelet abortion results in the all-bristle phenotype of *svslr1* mutants

To detail the morphological defects in *svslr1* spikelet formation, we used Scanning Electron Microscopy (SEM) and compared early developmental transitions of inflorescence primordia from A10.1 normal and *svslr1* plants (Fig. 1I-L, Fig. S6). In A10.1, the transition from the vegetative to reproductive meristem occurred around 10 days after sowing (DAS) (Fig. S6). Primary BMs were formed by 12 DAS, and secondary and higher order branching was initiated between 14 and 15 DAS (Fig. S6). In contrast, primary BMs were already well-formed in *svslr1* mutants by 10 DAS (Fig. S6), indicating that the reproductive transition happened earlier in *svslr1*. Developmental advancement was also observed during sequential production of secondary and tertiary axillary branches in *svslr1* inflorescence primordia (Fig. S6). In A10.1, the IM terminated in an SM followed by differentiation of SMs and bristles from the top of inflorescence and continuing basipetally around 16 DAS; bristles and spikelets were well formed by 18 DAS (Fig. 2I-J, Fig. S6). In *svslr1*, some BMs were already elongated by 14 DAS, before any visible sign of SM or bristle differentiation (Fig. 1K, S6). This was not observed in A10.1 at the comparable stage.

**Fig. 2.**
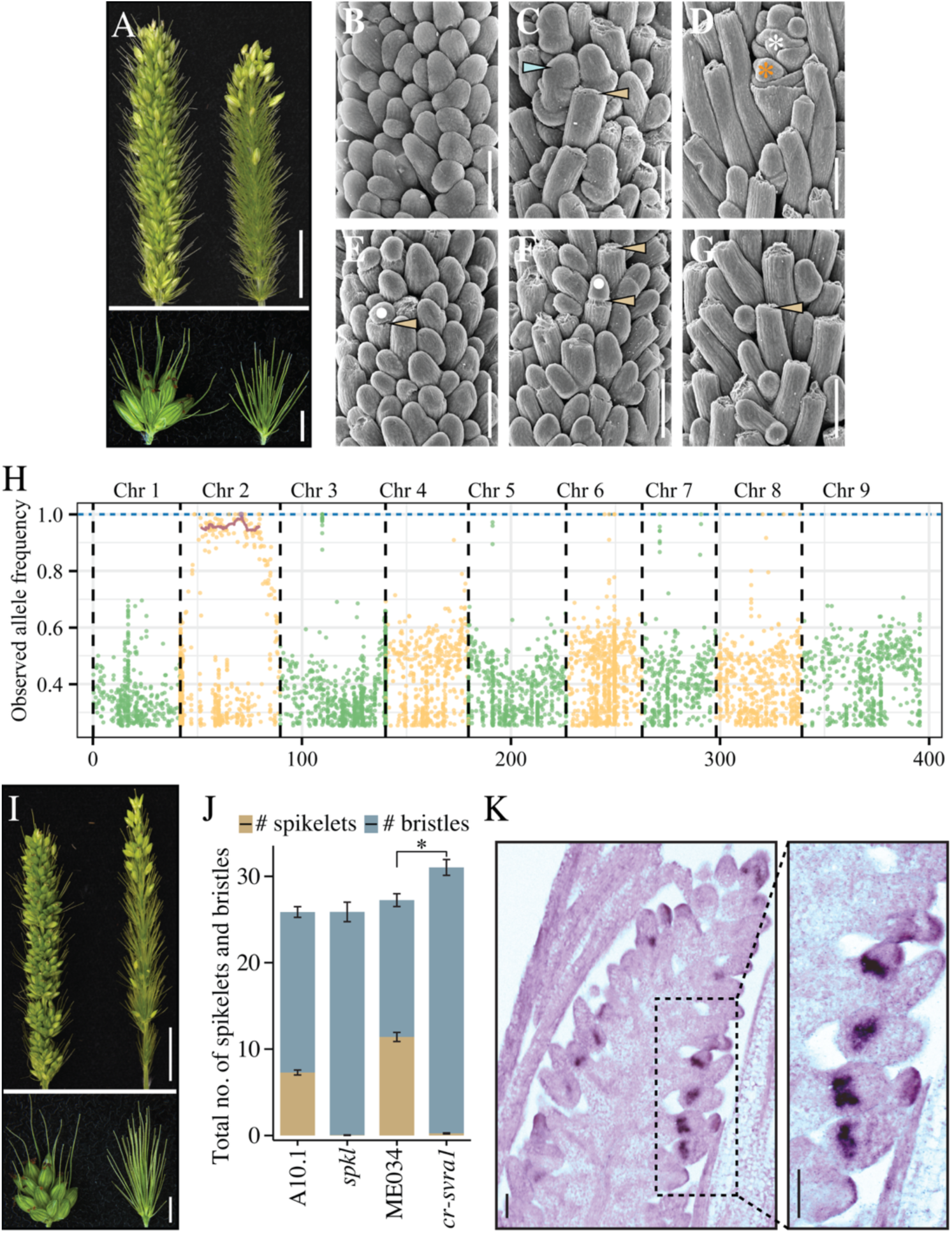
Characterization and genetic mapping of the *spikeletless* (*spkl)* mutant. (A) Panicles (top) and primary branch clusters (bottom) of A10.1 (left) and *spkl* (right) show that *spkl* mutants lack spikelets except for a few produced at the panicle tip. (B-G) SEM images of developing panicles displaying spikelet and bristle development in (B-D) A10.1 and (E-G) *spkl*. (B-D) In A10.1, (B) there were no apparent signs of spikelet meristems (SM) or bristle primordia at 16 DAS. (C) Bristles with severed meristem tips (gold arrow) and spikelet meristems (SMs, blue arrow) were visible at 17 DAS in A10.1. (D) Around 18 DAS in A10.1, SMs acquired determinate fate and differentiated into an upper fertile (white asterisk) and a lower aborted (orange asterisk) floret, and bristles were fully elongated. (E-G) In *spkl*, (E) a few BMs with an indented ring (gold arrow) around the meristem (white dot) were evident around 16 DAS. (F) By 17 DAS, some of the BMs (gold arrow) transitioned to form bristles with meristem tips broken and severed at the notch, while others (gold arrow) maintained the indented ring around the meristem (white dot). (G) SMs were still not visible and bristles (gold arrow) were fully developed in *spkl* by 18 DAS. (H) Genetic mapping of the *Spkl* locus using bulked segregant analysis. Allele frequencies (y-axis) of single nucleotide polymorphisms (SNPs, represented by green and yellow dots) were plotted across genomic positions (x-axis). DNA pooled from 34 *spkl* F₂ individuals (obtained from a *spkl* × A10.1 cross) was sequenced to identify the SNPs in *spkl* compared with A10.1 reference genome. An allele frequency = 1 indicates homozygosity. Red line highlights the homozygous SNPs (allele frequency ≥ 0.90) using a sliding window smooth curve over ten adjacent SNPs. Purple dot indicates the SNP that localized within *SvRa1* gene (I) CRISPR (CR)-cas9 based editing of *SvRa1* resulted in a ‘spikeletless’ phenotype. Panicles (top) and primary branch clusters (bottom) of *cr-svra1* (right) and parental genotype ME034 (left). The *cr-svra1* panicles lacked spikelets except for a few at the tip as in the *spkl* mutant. (J) Quantification of spikelets and bristles per primary branch cluster (shown in the lower part of A and I) of A10.1, *spkl,* ME034 and *cr-svra1.* Five primary branch clusters were taken from the middle of each panicle to count the average number of bristles/spikelets per cluster per panicle. For each panicle, averaged spikelet and bristle counts per cluster were summed; and a two-tailed t-test was used to compare *spkl* and *cr-svra1* (* *p* < 0.05, t-test) to their respective parental genotypes, A10.1 and ME034. Bars represent the mean +/- se of 28 plants/panicles. (K) *In-situ* hybridization showing *SvRa1* transcript localization in A10.1 panicles at 17 DAS. Localization of a *SvRa1*-specific antisense probe shows that *SvRa1* accumulated near the base of SMs and was absent from bristles. **Scale bars:** 1 cm (A and I, upper part), 2 mm (A and I, lower part), 100 μm (B-G), 50 μm (K).

Around 15 DAS, both SMs and bristles developed in *svslr1* mutants, however we observed initiation of an indented ring, a marker for BM to bristle transition (Yang et al. 2018), around the tip of SMs (Fig. 1L). The indented ring continued to sever the SM tips, leading to abortion of SMs in *svslr1* and leaving scars at the base of elongated bristles (Fig. S6). Counting fully developed terminal structures (spikelets and bristles) within a primary branch cluster of mature *svslr1* panicles indicated that SMs were aborted rather than converted to bristles and the number of bristles in *svslr1* was lower than in A10.1 (Fig. 1C, Fig. S6, Table S1). These results suggest that hyper-responsiveness to GA_3_ due to loss of function in DELLA likely results in enhanced elongation of differentiating axillary meristems, leading to spikelet abortion and long bristles in *svslr1*.

### Spikelets are converted to bristles in the *spikeletless* mutant

Compared to A10.1, *spkl* mutant panicles were smaller and showed increased primary branch density (Table S1). Within the mutant primary branch clusters that largely produced only bristles and no spikelets, bristles appeared to be paired (Fig. 2A, S1). In addition to the obvious panicle defects, *spkl* also showed several vegetative phenotypes including reduced plant height and increased tiller number (Fig. S7, Table S2).

Using SEM to compare early inflorescence development in *spkl* and A10.1 normal plants, we observed that the vegetative-to-reproductive transition and initial branching events were unaffected in *spkl* mutants (Fig. S8). In both A10.1 and *spkl*, primary BMs were formed by 13 DAS and secondary and higher order branching was initiated between 14 and 15 DAS. Around 16 DAS, the IMs of A10.1 and *spkl* transitioned to form the first SMs, which also marked the appearance of developmental differences between in *spkl* primordia (Fig. S8). In A10.1, most BMs were indistinguishable at 16 DAS (Fig. 2B) and differentiation of bristles and SMs was complete by 17 DAS; BMs set to become bristles had formed an indented ring around the meristem tip, which was ultimately lost (Fig. 2C, Fig. S8) and BMs that developed into SMs initiated glumes. At 18 DAS, the upper FMs formed perfect flowers with four whorls of floral organs and the lower FMs ceased development in A10.1 (Fig. 2D). In contrast, only short BMs or those with an indented ring around the meristem tip were observed at 16 DAS in *spkl* mutant primordia (Fig. 2E). Eventually, all BMs elongated and lost their meristem tip as they differentiated into bristles (Fig. 2F-G, Fig. S8). These results suggest that BMs poised to become spikelets in normal inflorescence development instead take on a bristle fate in *spkl* mutants. This phenomenon is directly opposite to what is observed in *bsl1* mutants, where bristles are homeotically converted to spikelets (Yang et al. 2018), and in stark contrast to the abortion of spikelet development observed in *svslr1* mutants.

### *Spkl* encodes the setaria functional ortholog of the maize RAMOSA 1 transcription factor

The *spkl* mutant was crossed to the parental A10.1 line and resulting F_2_ populations from selfed F_1_ individuals displayed the expected Mendelian 3:1 ratio for a single locus recessive allele (188:63; P (χ2, 1 d.f.) =0.97). We used bulk segregant analysis (BSA) to map the *Spkl* locus. We sequenced DNA pooled from 34 *spkl* mutant individuals segregating among F_2_ populations. Reads were mapped to the A10.1 reference genome (phytozome.jgi.doe.gov; v2.1, (Goodstein et al. 2012)) and single SNP sites were determined using Samtools mpileup (Version=3.5-0-g36282e4) (Supplemental dataset 1). BSA revealed a 10.2 Mb region (21740453-31964154) on chromosome 2 that showed high homozygosity in the *spkl* mutant pool (Fig. 2H). Within this interval, a non-synonymous, homozygous SNP was identified in *Sevir.2G209800*, the ortholog of maize *ramosa1* (*ra1*). We sequenced the *SvRa1* gene in *spkl* individuals and identified a 49 bp deletion in its only exon (Fig. S7). Genotyping validated that this deletion co-segregated with the ‘spikeletless’ phenotype (Fig. S7).

To confirm *SvRa1* as the gene responsible for the *spkl* phenotype, we generated a 101 bp deletion in the 5՛ end of the *SvRa1* exon in ME034 (Fig. S7). T_2_ plants that maintained the deletion were backcrossed to ME034 and then selfed to isolate Cas9-free, homozygous *cr-svra1* plants (Fig. S7), which showed a ‘spikeletless’ phenotype (Fig. 2I). Consistent with *spkl* phenotypes, *cr-svra1* plants also showed increased tillering compared to ME034 (Fig. S7, Table S3) and retained a few spikelets at the panicle tip. An allelic test between *spkl* and *cr-svra1* showed no complementation in the F_1_, confirming that *SvRa1* was responsible for the *spkl* phenotype (Fig. S9).

Since the SEM data indicated a homeotic conversion of spikelets to bristles, we examined the total number of bristles and spikelets within primary branch clusters from *spkl* and *cr-svra1* mutants and their parental lines (Fig. 2J). Five primary branch clusters were collected from the middle of the panicle and average counts of bristles and spikelets per cluster were recorded. Consistent with SEM data, both *spkl* and *cr-svra1* produced mostly all bristles, yet the aggregate of bristles and spikelets was like their parental control lines.

In maize, *ra1* is specifically expressed during the BM to SPM transition and transcripts localize to a boundary at the base of SPMs (Vollbrecht et al. 2005; Eveland et al. 2014). Based on published transcriptome data from developing setaria inflorescence primordia (Zhu et al. 2018), *SvRa1* transcripts accumulated beginning ∼14 DAS (prior to establishment of SM identity) and peaked at ∼16 DAS when most axillary meristems have differentiated into either a spikelet or a bristle (Fig. S8). We performed *in-situ* hybridization using a *SvRa1*-specific antisense probe to examine transcript accumulation in SMs and bristles of A10.1 inflorescence primordia at 17 DAS (Fig. 2K, S10). *SvRa1* transcripts were observed at the base of axillary meristems that had initiated development of SMs. We did not observe *SvRa1* accumulation in newly differentiated bristles. Compared with patterns of *Bsl1* and *SvBd1* transcript accumulation (Yang et al. 2018), *SvRa1* appeared to express in adjacent or overlapping domains.

### *SvRa1* acts upstream in the same genetic pathway as *Bsl1*

Since *spkl* and *bsl1* mutants appear to have directly opposite effects on bristle vs spikelet fate, we tested whether *SvRa1* and *Bsl1* act in the same genetic pathway. We made *spkl*;*bsl1* double mutants by crossing *spkl* to the strong *bsl1-1* allele (Fig. S11). Double mutant panicles looked like *bsl1* mutants with mostly spikelets produced (Fig. 3A). Like *bsl1*, pedicels of primary branches in the double mutant panicles were shorter (Fig. S11; (Yang et al. 2018)). SEM analysis of inflorescence development in *spkl;bsl1* double mutants showed a similar progression as was observed in *bsl1* mutants: 1) all higher-order BMs of the *spkl;bsl1* double mutant differentiated into spikelets; 2) some lower floral meristems developed normal florets rather than abort (Fig. 3B). This indicates that *SvRa1* and *Bsl1* function in the same pathway and *Bsl1* is epistatic to and downstream of *SvRa1*.

**Fig. 3.**
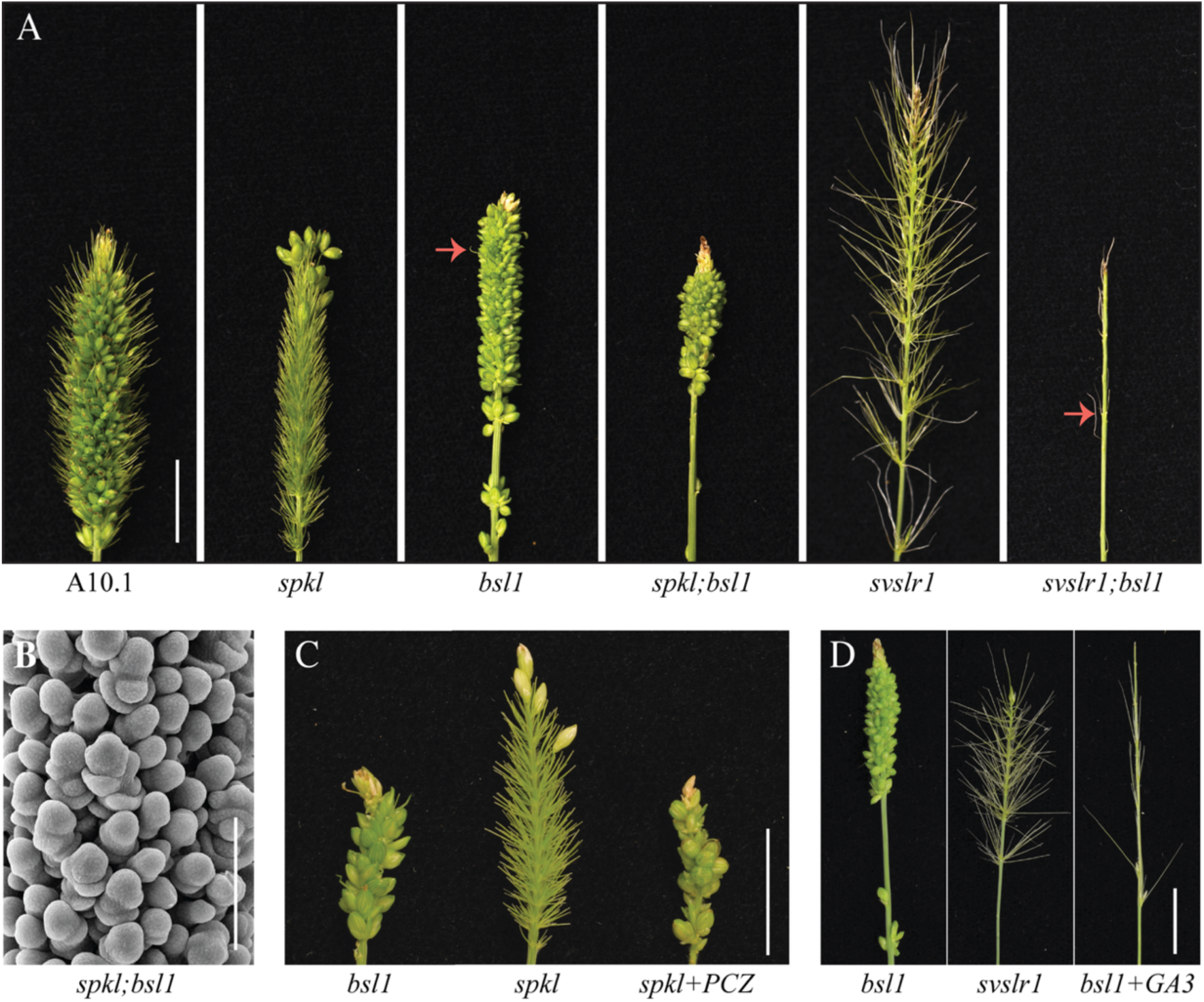
Genetic interactions between BR and GA genes regulate spikelet vs. bristle fate. (A) Genetic interactions between *bsl1*, *spkl* and *svslr1*. Representative panicles *of* A10.1, *spkl, bsl1, spkl;bsl1, svslr1 and svslr1; bsl1*. The double mutant *spkl;bsl1* looked like *bsl1* panicles and produced only spikelets. The *svslr;bsl1* panicles were mostly barren except for a few randomly arranged bristles (red arrow) that are typically produced in *bsl1* mutants. (B) Scanning Electron Microscopy of developing *spkl;bsl1* panicles showed formation of spikelet meristems and lack of bristle initiation. (C) Treatment of *spkl* mutants with the BR biosynthesis inhibitor propiconazole (PCZ) phenocopied the ‘bristleless’ trait of *spkl;bsl1*and *bsl1* panicles. Panicles from untreated *bsl1* (left), untreated *spkl* (middle) and PCZ-treated *spkl* (right) plants. (D) GA_3_ treatment of *bsl1* plants recapitulates the barren panicle phenotype of *slr1;bsl1*. Panicles from untreated *bsl1* (left), untreated *svslr* (middle) and GA_3_*-*treated *bsl1* (right). **Scale bars:** 1cm (A, C, D), 100 μm (B).

We previously showed that treating A10.1 plants with propiconazole (PCZ), a BR inhibitor (Hartwig et al. 2012; Sekimata et al. 2002), phenocopied the *bsl1* phenotype by converting bristles to spikelets (Yang et al. 2018). To test the effect of BR inhibition on *spkl* mutants, we applied 250 mM PCZ as a soil drench at 13 DAS (prior to differentiation of terminal BMs) and continued watering with PCZ for a week as previously described (Yang et al. 2018). Compared to non-treated mutant plants, treated *spkl* seedlings showed severe reduction in height (Fig. S11) and strikingly, the ‘spikeletless’ phenotype was converted to a ‘bristleless’ phenotype (Fig. 3C). This indicates that BR inhibition converts bristles to spikelets even in the absence of *SvRa1*.

### BR-mediated bristle vs spikelet fate is determined before a hyper response to GA aborts spikelets

So far, we showed that suppressing BRs by PCZ application converted bristles to spikelets (phenocopied *bsl1*) and that application of GA_3_ aborted spikelets and elongated bristles (phenocopied *svslr1*). Next, we examined the genetic interaction between *SvSlr1* and *Bsl1* by generating the *svslr1;bsl1* double mutant. The *svslr1;bsl1* plants were shorter than *svslr1* but taller than *bsl1* single mutants, and tillering was reduced compared to *bsl1* but increased when compared to *svslr1,* suggesting an additive role of GA and BR in controlling these traits (Fig. S11, Table S2). The *svslr1;bsl1* panicles were mostly barren with only a few elongated bristles (Fig. 3A). This phenotype can be explained by bristles being converted to spikelets in response to BR deficiency, which are then aborted through hyper-response to GA. Since *bsl1* (BR deficient) mutants typically produce a few randomly placed bristles, this is consistent with the few bristles formed in the double mutant. Treatment of *bsl1* plants with 10 mM GA_3_ applied at 14 DAS phenocopied the *svslr1;bsl1* double mutants (Fig. 3D).

We also tested the genetic interaction between *SvSlr1* and *SvRa1* by making the double mutant, *svslr1;spkl* (Fig. S11). Double mutant panicles were elongated and had elongated bristles like *svslr1*, but bristles were densely packed, suggesting that spikelets were converted to bristles as in *spkl* mutants rather than aborted. Unlike the *spkl* single mutant, we did not observe any spikelets retained at the panicle tip in *svslr1;spkl* likely due to spikelet abortion caused by the mutation in *SvSlr1*.

### Contrasting expression of meristem identity and determinacy regulators drive opposing phenotypes of *spkl* and *bsl1* mutants

To further identify factors involved in the bristle vs. spikelet fate decision, we compared transcriptomes of *spkl* and *bsl1* mutants with A10.1 normal inflorescences at three stages: 1) higher-order BM formation, 2) pre-fate specification and 3) early fate specification (Fig. S12). Differentially expressed (DE, adjusted *p* <0.05) sets of genes were identified for both mutants across the developmental stages (Table S4, S5). We hypothesized that genes and pathways core to the bristle vs. spikelet decision would show contrasting expression profiles in *spkl* and *bsl1* mutants; e.g., genes up-regulated in *spkl* and down-regulated in *bsl1* would promote bristle fate while genes with the opposite expression would promote spikelet identity. Comparing mutants to A10.1 at respective stages identified 232, 1048, and 7911 DE genes in *spkl* at stages 1, 2 and 3, respectively and 513, 1624, and 7240 DE genes in *bsl1*, respectively (Fig. S12).

Increasing numbers of DE genes observed during developmental progression indicates that molecular phenotypes of mutants begin to diverge around pre-fate specification (stage 2). At stage 2, *spkl* and *bsl1* shared 269 DE genes in common, including 81 genes that were up-regulated in *spkl* and down-regulated in *bsl1*, and 35 with the opposite profile (Fig. 4A, S12, Table S6).

**Fig. 4.**
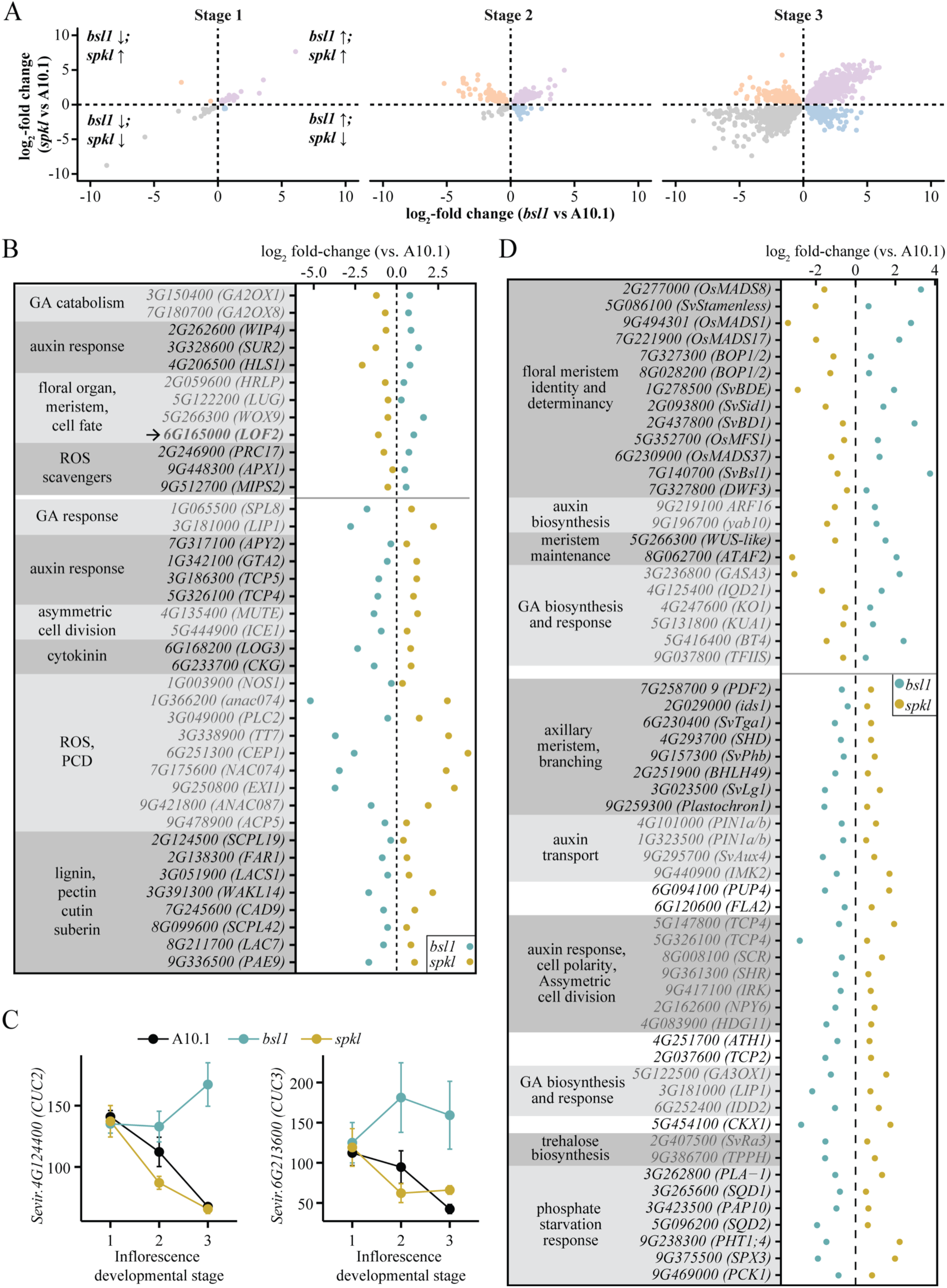
Transcriptome analyses of developing inflorescence primordia identified genes with contrasting expression in *bsl1*and *spkl*. (A) Scatterplots showing relative expression (log_2_-fold change compared to A10.1 controls) for genes that were differentially expressed (DE, *p*adj<0.05) in both *bsl1* and *spkl* compared to A10.1. Stage 1, 2, and 3 represent consecutive developmental stages of inflorescence as explained in Supplementary Figure S12. Arrows indicate up- (↑) and down-regulation (↓) relative to A10.1. (B) At stage 2 of inflorescence development, DE genes with opposite expression patterns in *bsl1* and *spkl* included orthologs of genes involved in auxin response, gibberellin (GA) catabolism and response, reactive oxygen species (ROS) metabolism, programmed cell death (PCD), floral organ development, and meristem and cell fate regulation. Black arrow marks an ortholog of the organ boundary gene, *LOF2*. Here, *Setaria virdis* gene identifiers (without prefix *Sevir*.) are noted along with gene names derived from *Arabidopsis thaliana* annotations; corresponding references are mentioned in Table S6. (C) Expression profiles of orthologs of *CUP-SHAPED COTYLEDON 2* (*CUC2*) and *CUC3* in developing panicles. (D) 59 selected DE genes implicated in processes related to meristem identity, determinacy and maintenance, auxin and GA biosynthesis and response, axillary meristem branching, auxin transport and asymmetric cell divisions, trehalose biosynthesis, and phosphate starvation response. These functional processes represent enriched Gene Ontology terms (details in Table S6) among gene sets that have contrasting expression in *bsl1* and *spkl* at stage 3. Gene identifiers are *Setaria virdis* gene IDs (without prefix *Sevir*.). Gene names from setaria and rice are indicated using the prefix *Sv* and *Os*, respectively. Maize and Arabidopsis gene names are indicated using lower- and upper-case letters, respectively.

There were few Gene Ontology (GO) terms enriched (adjusted p < 0.05) among these small sets of shared DE genes, and those that were did not provide clear functional clues; e.g., GO terms related to “chlorophyll catabolism” (GO:0015996) and “fatty acid derivative metabolic process” (GO:1901568) were overrepresented among genes up-regulated in *spkl* and down-regulated in *bsl1*) (Fig. S12, Table S7). However, analysis of individual genes and their orthologs in these DE gene sets identified regulators of stem cell fate and organ boundaries (Lee et al. 2009; Crawford et al. 2015) that were up-regulated in *bsl1* and down-regulated in *spkl*, including orthologs of *LOF2 (Sevir.6G165000*) and *WUSCHEL-related homeobox 9 (WOX9: Sevir.5G266300;* Fig. 4B, Table S6), consistent with setting of organ boundaries as SM identity initiates. Organ boundary genes, *CUC2* and *CUC3* (Hibara et al. 2006) also showed higher expression in *bsl1* than *spkl* beginning at stage 2 (Fig 4C).

In contrast, genes promoting programmed cell death (PCD) in floral tissues via accumulation of reactive oxygen species (ROS) (Gao et al. 2018; Zhang et al. 2014) were up-regulated in *spkl* and down-regulated in *bsl1*, while ROS scavengers (e.g., peroxidases; (Donahue et al. 2010; Xie et al. 2022; Davletova et al. 2005)) showed the opposite trend suggesting their role in tissue necrosis during bristle formation (i.e., formation of the indented ring). Genes up-regulated in *spkl* and down-regulated in *bsl1* also included regulators of cell polarization and genes involved in lignin, cutin, and suberin biosynthesis. Notably, GA catabolism genes (e.g., *GA2-oxidases*) and GA-responsive genes (Unte et al. 2003; Rombolá-Caldentey et al. 2014; Zhao et al. 2010) were down- and up-regulated in *spkl*, respectively, with opposite expression trends in *bsl1.* This indicates that GA response is enhanced in *spkl* (all bristles) and reduced in *bsl1* (all spikelets) during pre-fate specification (stage 2).

At stage 3, 2732 DE genes were common for *spkl* and *bsl1*, including 266 up-regulated in *spkl* and down-regulated in *bsl1*, and 268 with the opposite profile (Fig. 4A, S12). DE genes with contrasting expression profiles in the mutants were enriched for GO terms related to organ morphogenesis and auxin transport (Fig. S12, Table S7). Consistent with increased determinacy of terminal BMs in *bsl1* mutants, floral organ identity and meristem determinacy regulators were up-regulated in *bsl1* but suppressed in *spkl*, while genes known to promote branching and indeterminacy (e.g., *liguleless1, Phabulosa, Plastochron1,* (Lewis et al. 2014; McConnell et al. 2001; Sun et al. 2017)) were up-regulated in *spkl* and down-regulated in *bsl1* (Fig. 4D, Table S6). Genes involved in polar auxin transport (*SvAuxin4* (*SvAux4*) and *PIN-FORMED1* (*PIN1*)) and asymmetric cell divisions were also up-regulated in *spkl* with contrasting down-regulation in *bsl1* (Fig. 4D, S12, (Zhu et al. 2022)), potentially related to rapid cell growth and expansion during bristle formation. We found overrepresentation (*p* < 0.01) of TFs among genes with contrasting expression in *bsl1* and *spkl* at stage 3, suggesting that large-scale transcriptional rewiring underlies the opposing phenotypes of the mutants (Table S9). Among DE TF families, bHLH, SBP and TCPs were enriched among genes that were up-regulated in *spkl* and down-regulated in *bsl1* (Table S9, Fig. S13), suggesting that these TFs are negatively regulated by SvRA1 and positively by BR. Up-regulation of MADS-box TFs, known to regulate floral organ formation, in *bsl1* and their suppression in *spkl* is consistent with spikelet development.

### Features of the bristle vs. spikelet fate program are conserved in maize spikelet identity and determinacy

We isolated two loss-of-function alleles in maize *dwarf 11* (*d11*; *GRMZM2G107199*), the functional ortholog of *Bsl1*: 1) *d11-mu* was isolated from the UniformMu (W22 background) transposon population (McCarty et al. 2005) and 2) *d11-AcDs*, an Ac/Ds tagged allele, was generated using transposition in an Ac/Ds line (T43 background) with a Ds insertion (original stock) approximately 5.5kb downstream from *d11* (Ahern et al. 2009). The *d11-mu* allele contains a Mu insertion in the second intron of *d11* and *d11-AcDs* has Ds inserted in the second exon (Fig. S14). Compared to their respective genetic parents, W22 and T43, both *d11-mu* and *d11-AcDs* showed delayed flowering and semi-dwarf stature that was caused by shorter internodes (Fig. S14, Table S9). Tassel length and primary branch length in *d11-mu* were shorter than in W22 (Fig. 5A, S14, Table S9). The central tassel spike was shorter in both *d11-mu* and *d11-AcDs* compared to respective parental lines but tassel primary branch number was unaffected (Fig. S14, Table S9). Both alleles showed reduction in secondary branching compared to their respective parental lines (Fig. 5B-C, S14). In maize, the higher order branching produces paired spikelets: one pedicellate and one sessile. Interestingly, the length of pedicels bearing the pedicellate spikelet were severely reduced in both *d11* mutants as well as a significant reduction in spikelet size (Fig. 5D-F, S14, Table S9). These results indicate that D11-mediated accumulation of BR is important for branch elongation and secondary branching in maize tassels. An allelism test between *d11-mu* and *d11-AcDs* showed that the two mutants failed to complement and are allelic (Fig. S15). Since both alleles showed very similar phenotypes, we used *d11-mu* for further experiments.

**Fig. 5.**
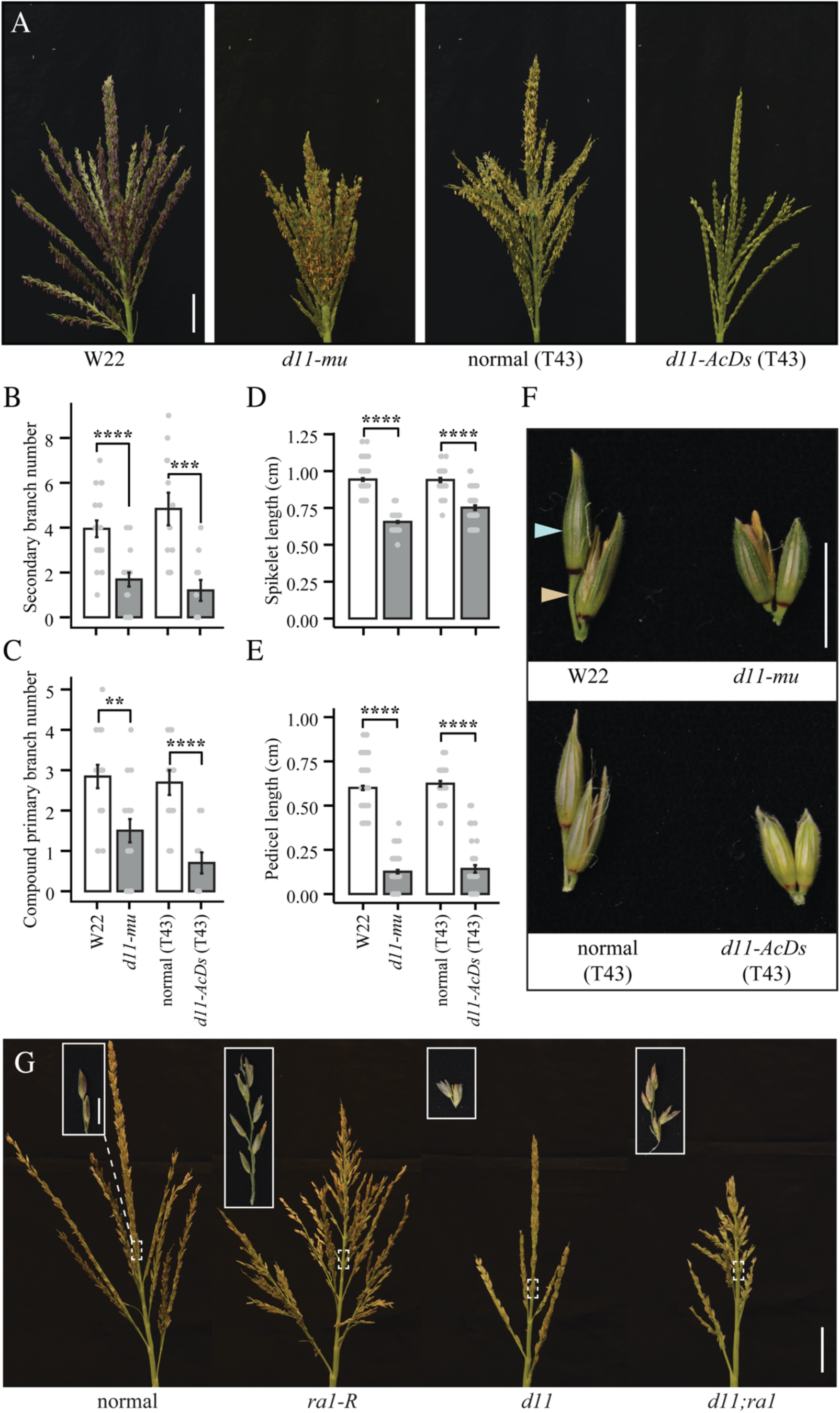
Phenotypic characterization of maize *dwarf 11* (*d11*) and genetic interaction with *ramosa 1.* (A-F) Characterization of the *d11-mu* allele (W22 background) and the *d11-AcDs* allele (T43 background). (A) Tassels from normal (W22) and *d11-mu* (W22 background) plants and segregating normal (T43) and *d11-AcDs* (T43) plants. (B-C) Both *d11* alleles showed a reduction in higher order branching. Both *d11-mu* and *d11-AcDs* showed (B) fewer secondary branches and (C) lower compound primary branches compared to corresponding non-mutant plants. (D-F) Spikelet pairs of *d11* alleles compared to corresponding non-mutants. (D) Pedicellate spikelet (blue arrow, F) and (E) pedicel (gold arrow, F) in *d11* mutants were shorter than in corresponding control genotypes. (G) Genetic interaction between *d11* and *ra1* in the B73 background. The *d11-mu* allele introgressed into B73 was crossed with the *ra1-R* reference allele in B73. Representative tassels from segregating normal sibs, *d11, ra1* and *d11;ra1* individuals are shown. Insets show the type of branch or spikelet-pairs that formed near the upper end of the long branch zone. Normal sibs and *d11* plants produced spikelet pairs, while *ra1* and *d11;ra1* showed spikelet multimers. **Scale bars:** 5 cm (A, G), 1 cm (F and inset in G). **Data:** mean ± SE; ** *p* < 0.01, *** *p* < 0.001, **** *p* < 0.0001

To test the genetic interaction between maize *d11* and *ra1,* we first introduced the *d11-mu* allele into B73 by backcrossing four times. We then crossed *d11-mu* (B73) with *ra1-R* (reference allele in B73). In controlled greenhouse conditions, *d11-mu* (B73) plant and tassel phenotypes were consistent with those of *d11-mu* (W22) (Fig. S16, Table S9) except for a reduction in tassel primary branch number, which was observed in *d11-mu* (B73), indicating the presence of genotype-specific modifiers that control BR-mediated tassel architecture. We next analyzed segregating individuals from the cross between *d11-mu* (B73) and *ra1* in the field (summers 2023 and 2024). The *d11;ra1* double mutant had a semi-dwarf stature similar to that of the *d11-mu* (B73) single mutant (Fig. S17). In normal control tassels, long branch formation abruptly shifts to spikelet pairs on the central spike. The *ra1-R* tassels showed a gradual shift from long branches to mixed-fate short branches to spikelet pairs, and exhibited increased primary and secondary branching compared to controls (Fig. 5G, S17, Table S10, (Vollbrecht et al. 2005)). In *d11;ra1* double mutant tassels, secondary branching was significantly lower than in *ra1* single mutants and the number of long primary branches was reduced to normal controls (Tukey HSD adjusted *p* < 0.05; Table S5, Fig. 5F, S17). Comparing lengths of the long branch, mixed-fate short branch and central spike zones from tassels of all genotypes showed that *d11;ra1* mutants had a short branch zone like what was observed in *ra1*. Similar to *ra1* tassels, *d11;ra1* produced multimers instead of spikelet pairs (Fig. 5G, Table S10).

A two-way ANOVA showed that genetic interactions between *d11* and *ra1* were significant for tassel traits including secondary branching and long branch formation, but not for mixed-fate short branches. Like *ra1, d11;ra1* double mutants also produced branched ears (Fig. S17, Table S10). It appears that *ra1* and *d11* functions intersect in controlling secondary branching in maize; the *d11* mutation suppresses increased primary and secondary branching caused by loss of functional *ra1*. However, *d11;ra1* double mutants retained the mixed-fate short branches and spikelet-multimers observed in *ra1* single mutants.

## Discussion

Our collection of setaria panicle mutants, *bsl1*, *spkl* and *svslr1*, when compared to normal controls with bristle-spikelet pairs, represent a developmental series from all spikelets to all bristles. Morphological and transcriptional analyses of these phenotypes, including interactions with the addition of GA or inhibition of BR, provided mechanistic clues to how the bristle vs. spikelet developmental program is controlled.

Double mutant analyses highlighted genetic interactions among the mutants, most notably that *SvRa1* acts upstream of *Bsl1* in the same genetic pathway. Loss of *SvRa1* function converts spikelets to bristles and loss of BRs results in all spikelets. Loss of both make all spikelets and therefore it is BR deficiency that is important for boundary formation and SM identity. The lack of bristle-spikelet pairs in either case, indicates that SvRA1 sets up the BR gradient, either directly or indirectly, between the bristle-spikelet paired meritems. In both maize and setaria, *Ra1* transcripts accumulate in a boundary domain adjacent to and at the base of the SM (Fig. 3K; (Strable et al. 2023)). The role of SvRA1 in maintaining determinacy of the spikelet seems analogous to its role in maize where RA1 promotes determinate growth of the SPM and maintains SM fate in a non-cell autonomous manner (Vollbrecht et al. 2005; Yang 2011; Whipple 2017).

Based on our collective data, we propose a model by which SvRA1 spatially restricts BR signals between adjacent paired BMs poised to become a spikelet-bristle pair, acting non-cell autonomously to maintain SM fate in one of the BMs and not the other (Fig. 6). In maize, RA1 binds to and modulates both GA biosynthesis and catabolism genes, likely regulating meristem determinacy through GA homeostasis (Eveland et al. 2014). Our transcriptome data from setaria showed that *GA2 oxidases* were up-regulated in BMs with spikelet fate. Therefore, we speculate that SvRA1 specifically promotes GA catabolism in SM primordia, while higher levels of GA in adjacent bristle primordia promote bristle outgrowth. A GA gradient between adjacent BMs may spatially restrict BR signaling via DELLA-mediated inhibition of BZR1 DNA binding activity (Bai et al. 2012). We hypothesize that in bristle primordia, higher GA activity degrades DELLA and releases BZR1 to suppress boundary genes and promote indeterminacy. This is supported by up-regulation of GA-responsive genes (*SPL8* and *LIPASE1* (*LIP1*)) and down-regulation of GA catabolic genes (*GA2 oxidases*) in *spkl* (bristle fate) with the opposite pattern in *bsl1* (SM fate). This suggests that the bristle fate is indeterminate fate, which is further supported by increased expression of genes known to mark indeterminacy in bristles (e.g., up-regulated in *spkl*). For example, *liguleless1* in maize is directly repressed by RA1 as tassels shift to making short branches (Eveland et al. 2014).

**Fig. 6.**
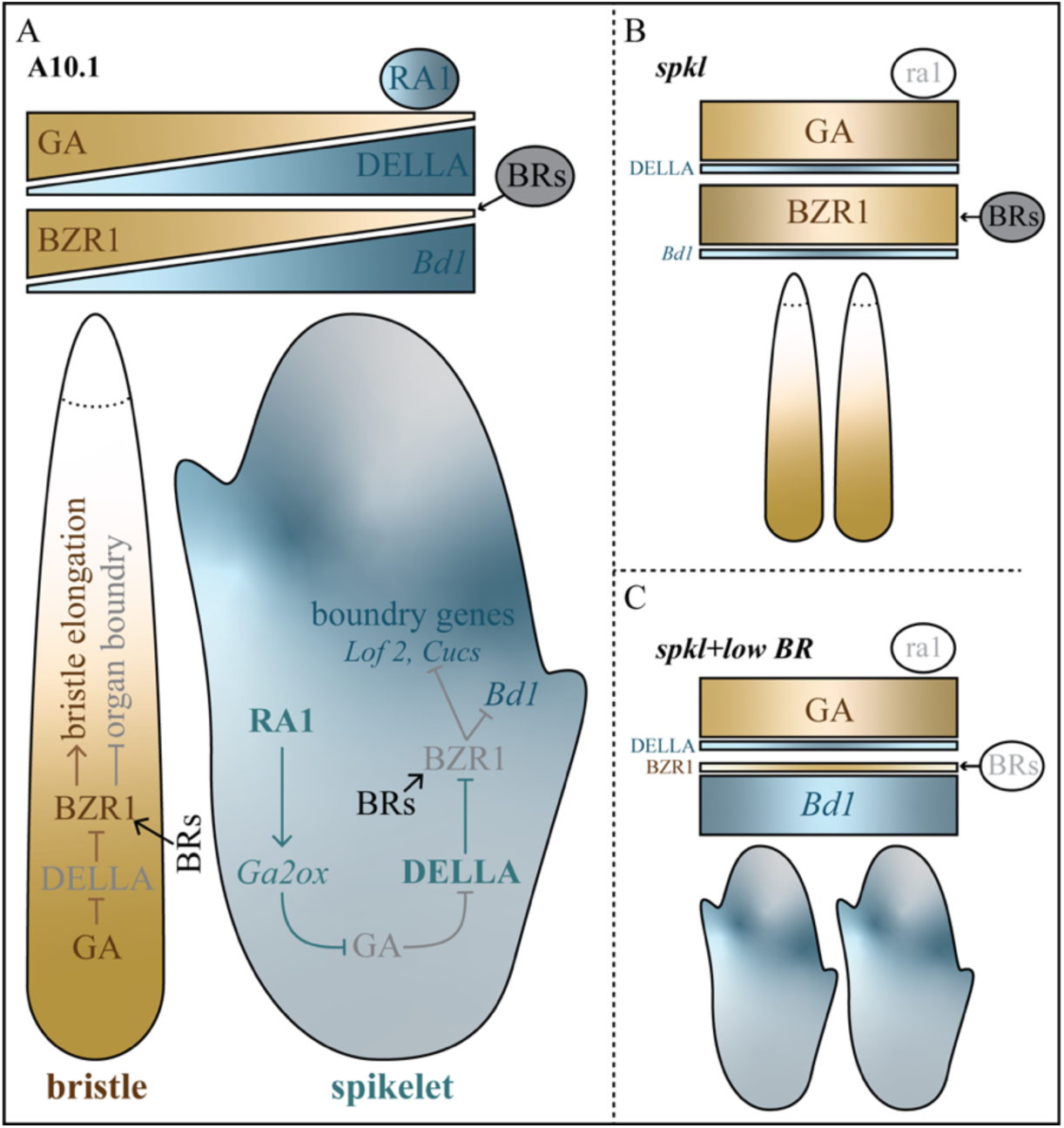
A proposed model for bristle vs. spikelet fate in paired branch meristems of developing setaria inflorescences. (A) RA1 enhances expression of the GA catabolism gene *GA2-oxidase* (*GA2OX*) in spikelet primordia resulting in a high-to-low GA gradient between bristle and spikelet primordia. In spikelets, lower GA levels would reduce DELLA degradation, thereby increasing DELLA accumulation. DELLA then inhibits DNA-binding activity of BZR1, which represses organ boundary genes. In bristles, higher GA concentrations would lead to DELLA degradation and therefore release of BZR1 from DELLA inhibition. Active BZR1 would then repress organ boundary genes and SM identity in bristles. BRs trigger the accumulation of the active form of BZR1. (B) In *spkl* mutants where RA1 is not functional, we propose that the GA gradient is abolished. This would result in lower DELLA levels and therefore release BZR1 from DELLA inhibition to promote the bristle program. (C) When BRs are reduced in *spkl* mutants through PCZ treatment or in *spkl;bsl1* double mutants, the active form of BZR1 would then be reduced, resulting in activation of organ boundary genes and initiation of SM identity.

BRs are implicated in placement of organ boundaries to precise domains during lateral organ formation through activation of BZR1. BR-mediated BZR1 expression suppresses boundary regulators (e.g., *CUCs* and *LOF2*) to restrict their expression to boundaries (Bell et al. 2012; Gendron et al. 2012). We found that in setaria, orthologs of *CUCs* and *LOF2* were up-regulated early in development (at the pre-fate stage) in SMs (*bsl1*), consistent with decreased BRs. Their lower expression in *spkl* (all bristles) supports an antagonistic role of BR (negative) and *SvRa1* (positive) in regulation of boundary genes (Fig. 4C). As expected, the SM identity gene *SvBd1* was also more highly expressed in *bsl1*, consistent with repression by BRs (Yang et al. 2018), resulting in SM identity and activation of FM identity genes. Genetic and BR-inhibitor assays support our model - reducing BRs restores SM identity in an *SvRa1* loss-of-function background (i.e., *spkl;bsl1* and PCZ-treated *spkl*) (Fig. 6).

In maize, SMs are paired so RA1 confers determinacy on the SPM (pair) in no apparent gradient since the pair is determinate. The bristle-spikelet pair in setaria may be an evolution of the spikelet pair in maize or it may have arisen independently with RA1 similarly co-opted for this role. The RA1 protein has several differences in the two species. RA1 from maize has two well-characterized EAR motifs towards the C-terminal end. In comparison, the SvRA1 protein has three amino acid differences in the C2H2 zinc finger domain but outside the invariant core, one amino acid change (Q169E) in the C-terminal EAR motif, and is missing the second EAR motif close to C2H2 zinc finger domain (Gallavotti et al. 2010, Strable et al. 2023). Interspecies expression of the *SvRa1* coding region in *ra1-R* mutants recovered only wild-type phenotypes when its expression was driven by the maize *ra1* promoter including an upstream region. Interestingly, this promoter region in maize includes conserved noncoding *cis* elements found exclusively in Andropogoneae species but not in setaria (Strable et al. 2023). Perhaps the proposed gradient that SvRA1 sets up across BM pairs is related to specific elements in its promoter or to interactions with a cell-specific regulator of *SvRa1* expression.

Based on the conversion model that was proposed for maize, the SPM forms an axillary SM and then converts into a second SM (Irish 1998). These two SMs form paired spikelets: one sessile and other pedicellate. Auxin inhibition assays and analysis of the *suppressor of sessile1* (*sos1*) mutant show that sessile spikelets develop from axillary SMs (Wu and McSteen 2007; Wu et al. 2009). Maize *sos1* is defective in auxin transport and forms single pedicellate spikelets (Wu et al. 2009). Perhaps the bristle-spikelet pair in setaria is morphologically analogous to a pedicellate-sessile spikelet pair in maize. Our maize *d11* mutants showed severe reduction in pedicel length of the pedicellate spikelet, which could be morphologically equivalent to altered bristle fate in setaria *bs11* (Yang et al. 2018). In setaria, mutants of auxin influx carriers (AUX proteins) including *sparse panicle1* (*spp1*; aka *svaux1*) and other multigene *svaux* mutants show altered spikelet vs. bristle fate. The *spp1/svaux1* mutant also frequently produces bristles with spikelets formed at the tip (Zhu et al. 2022) and likely this is due to the inability of the indented ring to completely sever the meristem. Auxin transporters including *PIN1* and *SvAux4* were upregulated in bristles (*spkl*) and suppressed in spikelets (*bsl1*), suggesting that auxin transport is involved in elongation of the bristle primordium and/or in formation of the indented ring below the meristem. It is likely that setaria bristle and spikelet development also follows a conversion model, where the BM itself converts to a bristle (indeterminate) and the SM generated immediately at its flank takes determinate fate and forms a sessile spikelet (Hodge and Doust 2017). Our transcriptome results provided limited yet novel evidence for this. For example, during the pre-fate stage, *DEVELOPMENT-RELATED PcG TARGET IN THE APEX4* (*DPA4*) was up-regulated in bristles and down-regulated in spikelets (Table S6). DPA4 is required to maintain meristem cell fate by suppressing *CUC2* expression (Nicolas et al. 2022).

Finally, we found that hyper-GA response caused SM abortion, but elongated bristles were retained. One possible explanation is that hyper-GA response selectively aborts SMs. Spikelet abortion due to GA_3_ treatment was reported in several maize accessions (Best and Dilkes 2022). Rice SPINDLY enhances the suppressive functions of DELLA, and *spindly* knockouts show panicle elongation and spikelet abortion (Shimada et al. 2006). The *spkl* mutant displays spikelets on the panicle tip and some of these spikelets have elongated pedicels. Hyper-GA response in a *spkl* background (in *spkl;svslr1*) enhances the ‘spikeletless’ phenotype of *spkl* and results in an all-bristle phenotype. It is likely that an unknown suppressor of the GA pathway that suppresses the *spkl/cr-svra1* phenotype at the panicle tip, is abolished by hyper-active GA response. It is notable that bristle count is also reduced in *svslr1*, which could indicate negative effects of GA response on higher order lateral branches in setaria panicles, similar to what has been observed for vegetative lateral branches in other species (Robil et al. 2025). Previous reports suggest that in some bristle grasses including setaria, some SMs are aborted early in development (Doust and Kellogg 2002). The spikelet to bristle ratio in setaria is lower than in domesticated foxtail millet (Doust et al. 2005), indicating selective breeding for more spikelets. Spikelet retention could be linked to an important trade-off in source-sink relations and perhaps GA mediates selective abortion. It is interesting to speculate how GA-mediated spikelet abortion could be involved in mediating source-sink relations during reproductive development, particularly in stressful environments (Wang et al. 2024).

## Material and Methods

### Plant Materials, Growth Conditions and Phenotyping

The *spkl* and *svslr1* mutant alleles were isolated from NMU-mutagenized M2 population of *Setaria viridis* (Huang et al. 2017). Setaria plants were grown in the greenhouse under long-day conditions with 28/22 ± 2°C (day/night), 16 h light/8 h dark, and 50% relative humidity at the Danforth Center’s Plant Growth Facility. For both mutants, seeds from F_3_ or later generations derived from backcrosses between mutant allele and the parental genotype A10.1 were used for phenotyping. Quantitative data for plant and panicle traits were collected right after flowering and pairwise t-tests were used to compare genotypes. For SEM, RNA-seq, *in situ* hybridization, and chemical (PCZ or GA_3_) treatments, plants were grown inside high-light growth chambers (31°C/22°C [day/night], 12 h light/12 h dark, 50% relative humidity).

The maize *d11-mu* (W22) allele (mu1090290, stock ID: UFMu-12395) from the UniformMu (W22) transposon population (Settles et al. 2007) and the Ac/Ds line (133822470, stock ID: AcDs-00008, background: T43) were obtained from Maize Genetics COOP Stock Center (http://maizecoop.cropsci.uiuc.edu/). Development of *d11-AcDs* (T43) followed the protocol described by (Ahern et al. 2009). The *d11* single mutants and corresponding control plants were grown in a greenhouse (28/22 ± 2°C with 14 hour day length and 50% relative humidity) to collect plant stature and tassel branching data. A t-test was used to compare mutants with corresponding control plants.

For testing genetic interaction between *d11* and *ra1*, segregating individuals derived from cross between *d11-mu* (B73) and *ra1-R* (B73 background) were grown at the Danforth Center’s Field Research Site in Saint Charles, MO during the summers (June-August) of 2023 and 2024. Quantitative measurements for plant stature and tassel branching traits were collected right after flowering. Two-way ANOVA was used for each tassel trait to test statistical interaction between factors: *d11* and *ra1;* and post-hoc multiple comparisons were performed using tukey-HSD.

### Bulked Segregant Analysis

The *spkl* allele from a mutagenized A10.1 population was backcrossed to A10.1 and F_1_ plants were selfed to make segregating F_2_ families. DNA extracted from 34 F_2_ *spkl* individuals was pooled to prepare a DNA library using the NEBNext Ultra DNA Library Prep Kit for Illumina (NEB), with inserts selected for sizes between 500 and 600 bp. The library was sequenced using a 150-bp paired-end design on the Illumina Hi-Seq 4000 platform at the Department of Energy Joint Genome Institute (JGI). Reads were mapped against the setaria reference genome v2 using Bowtie 2 (version 2.5.4) with default parameters and the --dovetail option. SNPs were called using Samtools mpileup (version 1.6) and filtered with VarScan (v2.4.6) using the options --min-reads 2, --min-coverage 5, and --min-avg-qual 15 (Klein et al. 2018). SNPs within repetitive regions, non-canonical NMU-induced mutations, with high coverage (greater than 100), and allele frequencies below 0.25 were removed from the analysis using R. Homozygous SNPs with allele frequency ≥ 0.90 on chromosome 2 were highlighted using a sliding window smooth curve of ten adjacent SNPs.

### Scanning Electron Microscopy Analysis

Inflorescence primordia were hand-dissected from young A10.1, *spkl*, *svslr1* and *spkl;bsl1* plants. The samples were fixed in 4% paraformaldehyde and dehydrated according to the protocol outlined by (Hodge and Kellogg 2014). Following dehydration, the samples underwent critical point drying using a Tousimis Samdri-780a and were subsequently imaged by a Hitachi S2600 SEM at Washington University’s Central Institute of the Deaf. The SEM images were converted to black and white scale and level adjustment layer was applied to adjust the contrast using Photoshop.

### *In situ* hybridization

Setaria inflorescence primordia were fixed, embedded, and sectioned as described previously (Yang et al. 2018). Digoxigenin-UTP-labeled *SvRa1* sense and antisense probes were synthesized by *in vitro* transcription of fragments of *SvRa1* cDNA by DIG RNA labeling kit (SP6/T7; Roche). The probes were used to detect *SvRa1* transcripts in sections via hybridization as described in (Yang et al. 2018). The Digoxigenin signal was detected overnight using nitroblue tetrazolium and 5-bromo-4-chloro-3-indolyl phosphate (Roche) following manufacturer’s instructions. Images of sections were collected using a NiKon NiE Upright microscope. The images were processed using Photoshop to correct white balance.

### Phylogenetic Analysis

The coding sequences of *SLR-*like genes from Arabidopsis, setaria, foxtail millet, maize, sorghum, rice, barley and wheat were retrieved from Phytozome (phytozome. jgi.doe.gov; Supplemental Dataset S2, (Goodstein et al. 2012)). The coding sequences were aligned using ClustalW. A maximum likelihood phylogenetic tree was constructed with bootstrapping (1,000 iterations) in MEGA7 (Kumar et al. 2016). The evolutionary history was inferred using maximum likelihood method and Tamura-Nei model (Tamura and Nei 1993).

### RNA-Seq Library Construction, Sequencing, and Analysis

For transcriptome analysis, inflorescence primordia from A10.1, *spkl* and *bsl1* seedlings were collected at three consecutive developmental stages. These developmental stages correspond to A10.1 inflorescence primordia collected at 13 (stage 1), 14 (stage 2), and 16 (stage 3) DAS. To account for the developmental progression of mutants, *spkl* primordia were sampled at 12, 14, and 15 DAS, while *bsl1* primordia were sampled at 15, 16, and 17 DAS. For each developmental stage, 4 biological replicates per genotype were collected. Each biological replicate contains hand-dissected inflorescence primordia pooled from multiple seedlings.

RNA extraction was performed using the PicoPure RNA isolation kit (Thermo Fisher Scientific). RNA Poly-A+ RNA-seq libraries prepared from 1μg of total RNA using the NEBNext Ultra Directional RNA Library Prep Kit (Illumina, San Diego, CA, USA) were size-selected for 200-bp inserts and quantified by an Agilent bioanalyzer using a DNA 1,000 chip. The RNA-seq libraries were processed on an Illumina HiSeq 4000 platform at Novogene with a 150-bp paired-end sequencing design and yielded 21-34 million reads per data point. Read quality checks and processing were conducted using the wrapper tool Trim Galore (version 0.4.4_dev) with the parameters “–length 100 –trim-n –illumine.” The cleaned reads were mapped to the *S. viridis* transcriptome (Sviridis_500_version 2.1; Phytozome version 12.1, (Goodstein et al. 2012)) using Salmon (0.13.1) with the parameters “–validateMappings –numBootstraps 30,” based on an index generated by primary transcripts (n = 38,209). Two samples with poor correlation among replicates (pearson correlation cut-off > 0.86) were excluded from downstream analysis. Gene normalized expression levels (transcript per kilobase million [TPM]; Table S4) and the count matrix for downstream analyses were obtained from Salmon output files. Salmon output files were imported in R using the Bioconductor package tximport (Soneson et al. 2015).

DE analysis was performed using the Bioconductor package *DESeq2* (version 1.22.2) with default parameters for the Wald test. Benjamini–Hochberg method was used for multiple testing correction (adjusted *p* <0.05) to identify DE genes.

The *S. viridis* GO annotations were generated using the GOMAP (Wimalanathan and Lawrence-Dill 2021) pipeline as described previously (Yang et al. 2021). The GO annotation package was prepared using ‘makeOrgPackagè function in ‘AnnotationForgè package in R. GO term enrichment analysis was performed using ‘enrichGÒ function of ‘clusterprofiler’ package. Functional annotation files containing Arabidopsis and rice orthologs of *S. viridis* (version 2.1) genes were obtained from PhytozomeV12 (Goodstein et al. 2012). Maize orthologs (RefGen V4) where *S. viridis* gene identifiers were extracted using Phytozome-Biomart (Goodstein et al. 2012). Information on maize classical genes was obtained from MaizeGDB (https://www.maizegdb.org, (Woodhouse et al. 2021)) using maize RefGen V4 identifiers.

### Identification and Functional Classifications of Gene Candidates

For each developmental stage, genes with DE in both *spkl* and *bsl1* were compared to identify four sets of DE genes: 1) up-regulated in *spkl* and down-regulated in *bsl1*, 2) down-regulated in *spkl* and up-regulated in *bsl1*, 3) up-regulated in both *spkl* and *bsl1*, 4) down-regulated in both *spkl* and *bsl1*. For stages 2 and 3, GO enrichment analysis for all four sets of DE genes was performed as described above. For DE genes with contrasting expression at stage 2, each *S. viridis* gene identifier and their Arabidopsis (TAIR ID), maize, and rice homologs were searched in the literature for potential functions. References for these genes are cited in Table S6. TF family information was obtained using the prediction tool (https://planttfdb.gao-lab.org/prediction.php) on plantTFDB (https://planttfdb.gao-lab.org, (Jin et al. 2017)). The coding sequences (Sviridis_500_v2.1.cds_primaryTranscriptOnly.fa.gz) obtained from PhytozomeV12 (Goodstein et al. 2012) were used as input for predicting TF families. Hypergeometric test in R was used to test the overrepresentation of TFs or TF families. The formula and code used for TF enrichment analysis are detailed in Table S8.

### CRISPR/Cas9 gene editing

Genomic sequences for *SvRa1* (*Sevir.2G209800*) and *SvSlr1* (*Sevir.9G121800*) were obtained from Phytozome (*S. viridis* version 2.1). To minimize off-targets, guide(g) RNAs were designed using CRISPR-P version 2.0 (Liu et al. 2017). Two gRNAs were designed to target the 5’ end of *SvRa1* exon. For *SvSlr1*, two guides targeting DELLA domain and third guide targeting immediately downstream of DELLA domain were designed. Only one guide, however, resulted in 1bp deletion in *cr-slr1* downstream of DELLA domain. Two separate gRNA constructs for *SvRa1* and *SvSlr1* were prepared using a plant genome engineering toolkit (Čermák et al. 2017) as described previously (Yang et al. 2021). Briefly, gRNAs were combined into a level 0 construct, which was then inserted into a plant transformation vector. PCR-amplified fragments from pMOD_B_2303 were merged via golden-gate cloning with T7 ligase and restriction enzymes SapI/BsmBI, reinserting them into the pMOD_B_2303 backbone to express the gRNAs from the CmYLCV promoter, each flanked by a tRNA. This guide construct, pMOD_A1110 module containing *ZmUbi1* promoter driven Cas9 (wheat codon-optimized) and pMOD_C_0000 module were combined in a subsequent golden-gate cloning reaction with T4 ligase and AarI restriction enzyme into the pTRANS_250d plant transformation backbone. This final binary vector was transformed into *Agrobacterium tumefaciens* strain AGL1, which was then used for callus transformation of *S. viridis* accession ME034 at the DDPSC Tissue Culture facility. T_0_ plantlets were genotyped for the presence of the selectable marker, hygromycin phosphotransferase (Hpt) to validate transgenic individuals. Individual T_1_ plants with possible mutant phenotypes were selected and screened for mutations in targeted regions by Sanger sequencing/genotyping. These T_1_ mutants were self-pollinated to obtain T_2_ progeny. Primers used for vector construction and genotyping are listed in Supplemental Table S11.

## Acknowledgments

This work was funded by the National Science Foundation (NSF) Plant Genome Research Program award #1733606 and the NSF IOS Developmental Mechanisms award #2440006. The authors thank Dr. Jose Dinneny (Stanford University) for identifying and providing the *svslr1* panicle phenotype in the setaria NMU population. Very special thanks to Ms. Zhonghui Wang and Ms. Jessica Helms for support in planting, harvesting and genotyping, and Ms. Taylor Guiot for with tassel collections. The authors also thank the Joint Genome Institute (JGI) for sequencing. The work conducted by the U.S. Department of Energy JGI was supported by the Office of Science of the U.S. Department of Energy under Contract No. DE-AC02-05CH11231. Also, special thanks to personnel from the Danforth Plant Science Center Plant Growth Facility, Advanced Bioimaging Lab, the Field Research Site, and Plant Transformation Facility.

## Author Contributions

JY isolated and characterized setaria mutant alleles and double mutants, mapped the *Spkl* locus, and performed genetic crosses in maize. JS performed follow-up analyses on setaria mutants and characterized maize double mutants. JY performed SEM and *in situ* hybridization. JY performed the RNA-seq experiment and JS analyzed the data. MB made the gene edited lines. EB analyzed the BSA experiment. JS generated the figures and assembled supplementary information. JS developed the model with input from JY and ALE. JS, JY and ALE conceptualized the research and wrote the paper. All authors reviewed the final manuscript.

## Supplementary Information

### Supplementary Figures

**Fig. S1.** Setaria mutants with defects in spikelet or bristle formation.

**Fig. S2.** Tillering phenotype and CAPS marker based genotyping of *svslr1* mutant.

**Fig. S3.** Gene model of *SvSlr1* showing the location and sequence of the guide RNA used to generate edits in the *cr-slr1* mutant.

**Fig. S4.** Whole plant morphology of untreated A10.1, GA_3_-treated A10.1 and untreated *svslr1*.

**Fig. S5.** Expression of the *SvSlr1* gene in developing setaria inflorescences (based on data from Zhu et al., 2018).

**Fig. S6.** Scanning electron microscopy images showing developmental progression of A10.1 and *svslr1* inflorescence primordia.

**Fig. S7.** Whole plant morphologies of *spkl* and edited *cr-svra1* and gene model of *SvRa1* showing locations of deletions in *spkl* and *cr-svra1*.

**Fig. S8.** Scanning electron microscopy images showing developmental progression of A10.1 and *spkl* inflorescence primordia.

**Fig. S9.** Plants, panicles and genotyping of F1s obtained from genetic cross between *spkl* and *cr-svra1*.

**Fig. S10.** *In-situ* hybridization with the sense *SvRa1* control probe.

**Fig. S11.** Comparing whole plant and panicle morphologies of A10.1, single mutants (*bsl1, spkl* and *svslr1*) and double mutants (*spkl;bsl1, svslr1;bsl1, svslr1;spkl*); and effect of PCZ treatment on whole plant morphology of *spkl*.

**Fig. S12.** Schematic representation of RNA-seq analysis comparing transcriptomes of A10.1, *spkl* and *bsl1* inflorescence primordia and GO term enrichment analysis of DE gene sets

**Fig. S13.** Expression profiles of selected TF families in *bsl1*, A10.1, and *spkl* inflorescence primordia.

**Fig. S14.** Whole plant and tassel phenotypes of maize *dwarf11* (*d11*), *Bsl1* ortholog, alleles: *d11-mu* and *d11-AcDs*.

**Fig. S15.** Phenotypic characterization of F1s obtained from genetic cross between two alleles of *d11*: *d11-mu* and *d11-AcDs*.

**Fig. S16.** Phenotypic characterization of *d11-mu* allele introgressed into B73 background.

**Fig. S17.** Plant, tassel and ear phenotypes of normal, *d11-mu*, *ra1-R* and *d11-mu;ra1-R* plants obtained from genetic cross between *d11-mu* (B73 background) and the *ra1* reference allele (*ra1-R*; B73 background).

## Supplementary Tables

**Table S1.** Measurements of panicle traits for A10.1, *spkl, svslr1,* ME034 and cr-*svra1*.

**Table S2.** Plant height and tiller count data for A10.1, *spkl, svslr1, bsl1, svslr1;bsl1* and *spkl;bsl1* collected at the flowering stage.

**Table S3.** Plant height and tiller count data for parental genotype ME034 and *cr-svra1*.

**Table S4.** RNA-seq based transcript abundance (Transcripts Per Million, TPM) for all *Setaria viridis* (*Sv*, V2.1) genes in A10.1, *bsl1*, and *spkl* inflorescence primordia across the three developmental stages.

**Table S5.** DE genes in *bsl1* and *spkl* compared to parental genotype A10.1.

**Table S6.** DE genes with contrasting expression changes (log_2_-fold change relative to A10.1) in *bsl1* and *spkl* mutants.

**Table S7.** Gene Ontology terms enriched among gene sets that are DE in both *bsl1* and *spkl* mutants.

**Table S8.** Enrichment of TFs and TF families among gene sets that are DE in both mutants at stage 3.

**Table S9.** Plant and tassel traits for *d11* mutants and corresponding controls.

**Table S10.** Phenotypic analysis of *ra1, d11, d11;ra1* and normal plants obtained from crosses between *ra1* and *d11*.

**Table S11.** Primers used in the study.

