## Supplementary Information for "A core genetic mechanism integrates growth hormone signals to control meristem fate and inflorescence architecture in setaria and maize"

**Supplementary Data**

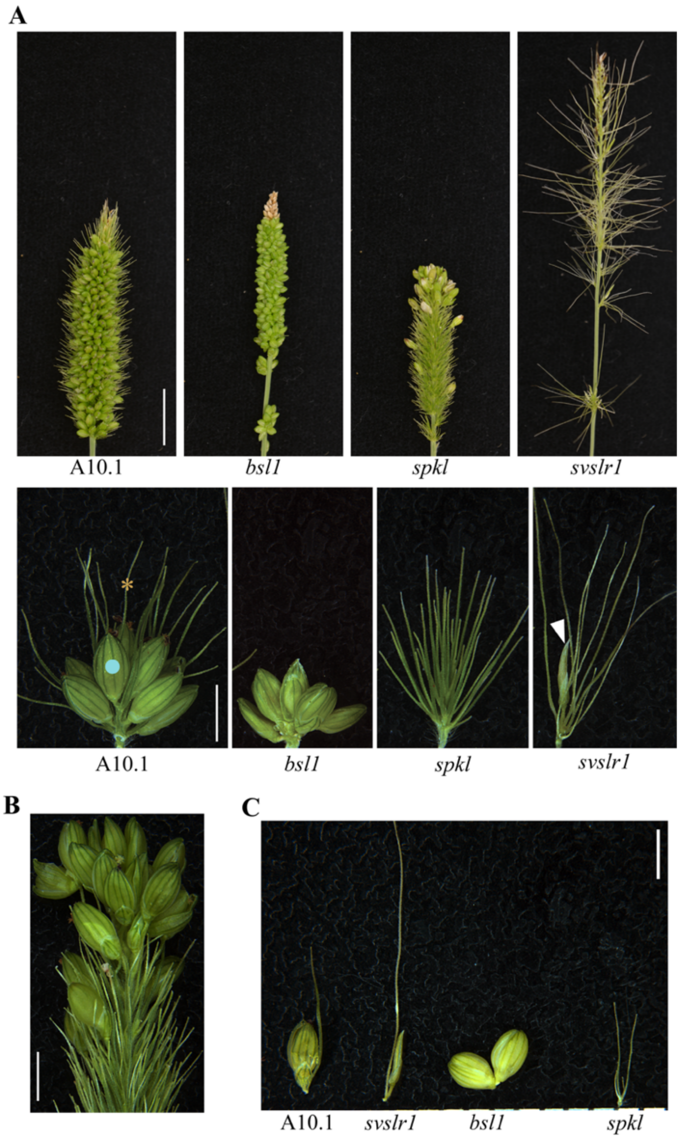


**Figure S1. Mutants showing defects in spikelet (blue dot) and bristle (gold asterisk) formation were isolated from N-nitroso-N-methylurea (NMU)-mutagenized population of *Setaria viridis* (parental accession A10.1).** (A) Panicles (top) and primary branch clusters (bottom) of A10.1, *bsl1* (*bsl1-1* allele from Yang et al. 2018), *spkl* and *svslr1*. (B) Image of *spkl* panicle tip showing a few spikelets are formed on otherwise spikeletless panicle. (C) Presumably paired spikelet-bristle in A10.1 and *svslr1*; spikelet-spikelet in *bsl1;* and bristle-bristle in *spkl*. White arrow: aborted spikelet in *svslr1*.

**Scale bars:** 1 cm (A-top), 2 mm (A-bottom, B, C).

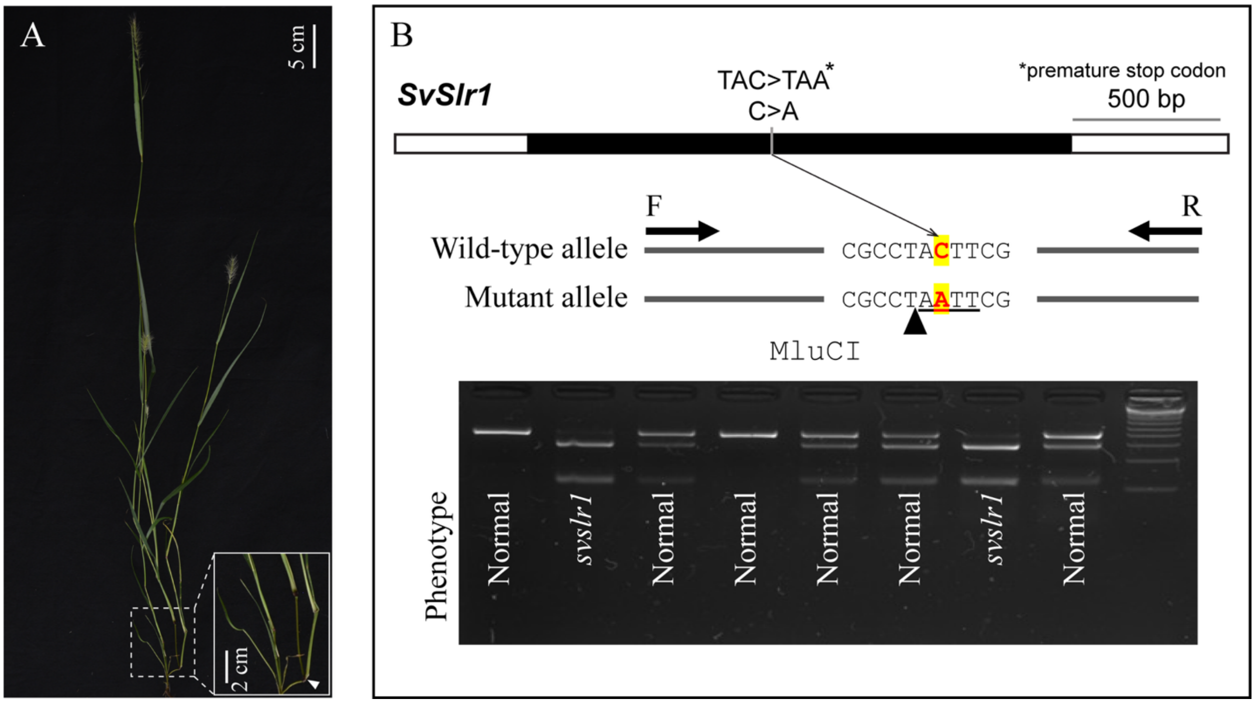


**Figure S2. Tillering phenotype and CAPS marker based genotyping of *svslr1* mutant**

(A) Plant morphology of *svslr1* with inset showing secondary tillers on nodes of primary tillers of *svslr1* mutant. (B) *SvSlr1* gene model (top) showing location of C>A transition in *svslr1* mutant. Genotyping using cleaved amplified polymorphic sequences (CAPS) marker with MluCI cut site showing plants with *svslr1* phenotype are homozygous for C>A.

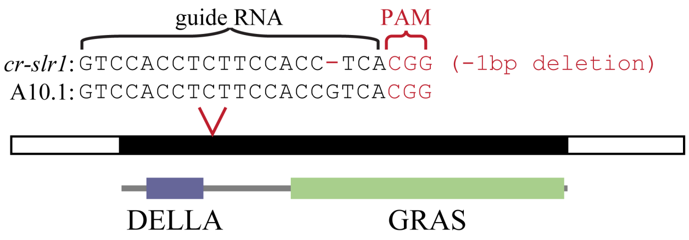


**Figure S3.** Gene model of *SvSlr1* showing the location and sequence of the guide RNA used to generate edits in the *cr-slr1* mutant. PAM: Protospacer Adjacent Motif. DELLA and GRASS are protein domains.

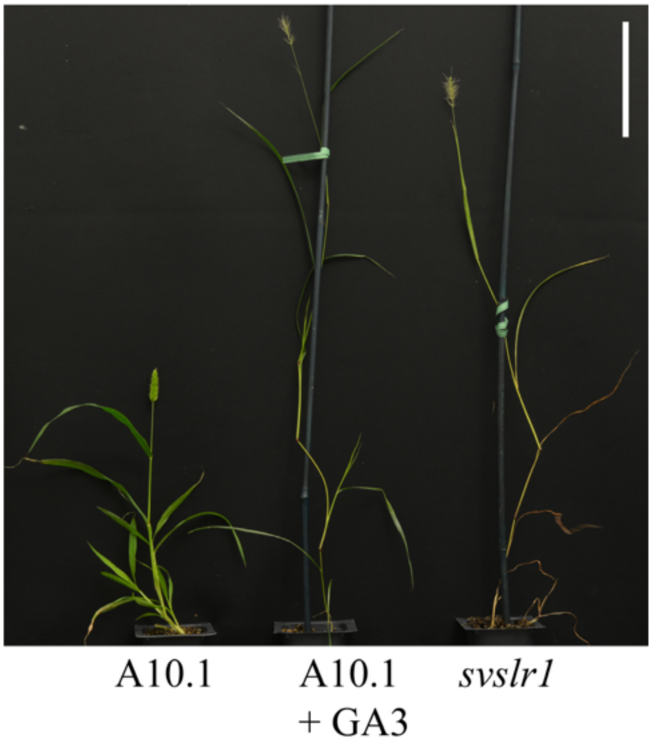


**Figure S4.** Whole plant morphology of untreated A10.1, $GA_3$-treated A10.1 and untreated *svslr1*.

**Scale bar:** 10 cm.

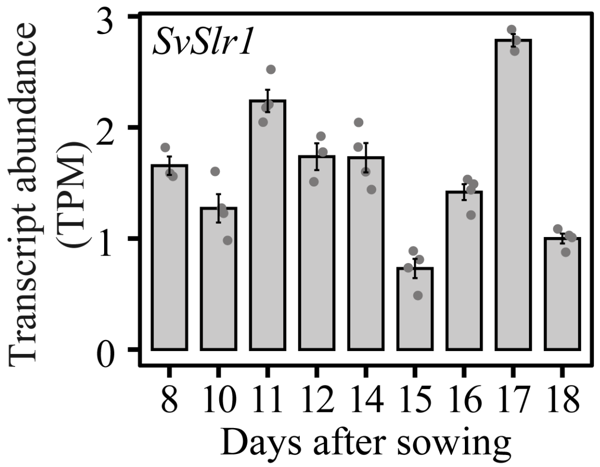


**Figure S5.** Expression of the *SvSlr1* gene in developing setaria inflorescences (based on data from Zhu et al., 2018.

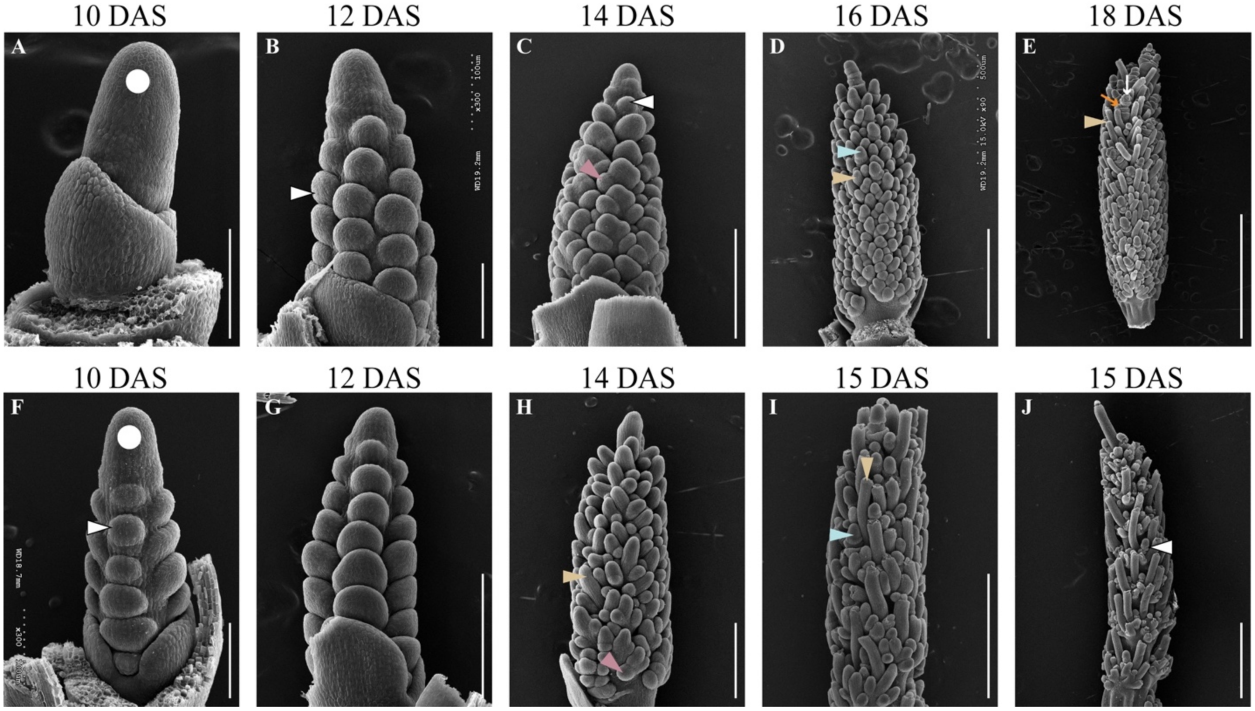


**Figure S6. Scanning electron microscopy images showing developmental progression of A10.1 and *svslr1* inflorescence primordia.**

(A-E) In A10.1, (A) inflorescence meristem (IM, white dot), (B) primary branch meristems (BMs, white arrows) (C) and higher order BMs (pink arrows) were visible around 10, 12 and 14 days after sowing (DAS), respectively. (D) IM (white dot) transitioned to terminal spikelet, and differentiation of spikelet meristem (SM, blue arrow) and bristle (gold arrow) was apparent by 16 DAS in A10.1. (E) Bristles with meristem broken off at indented ring (gold arrow) were fully developed by 18 DAS in A10.1. SM transitioned to floral meristem, resulting in formation of upper fertile floret (white arrow) and lower aborted floret (orange arrow).

(F-I) Compared to A10.1, *svslr1* showed developmental advancement. (F-G) primary BMs (white arrow) were already formed by 10 DAS. (H) IM transitioned to the determinate fate by 14 DAS and most BMs appeared elongated (gold arrow) in *svslr1.* (I) SMs (blue arrow) and full developed bristle (blue arrow) were already formed by 15 DAS in *svslr1*. (J) SMs (blue arrow) aborted before differentiation in *svslr1.*

**Scale bars:** 100 µm (A-B, F-G), 250 µm (C, H), 500 µm (D, I), 1 mm (E, J).

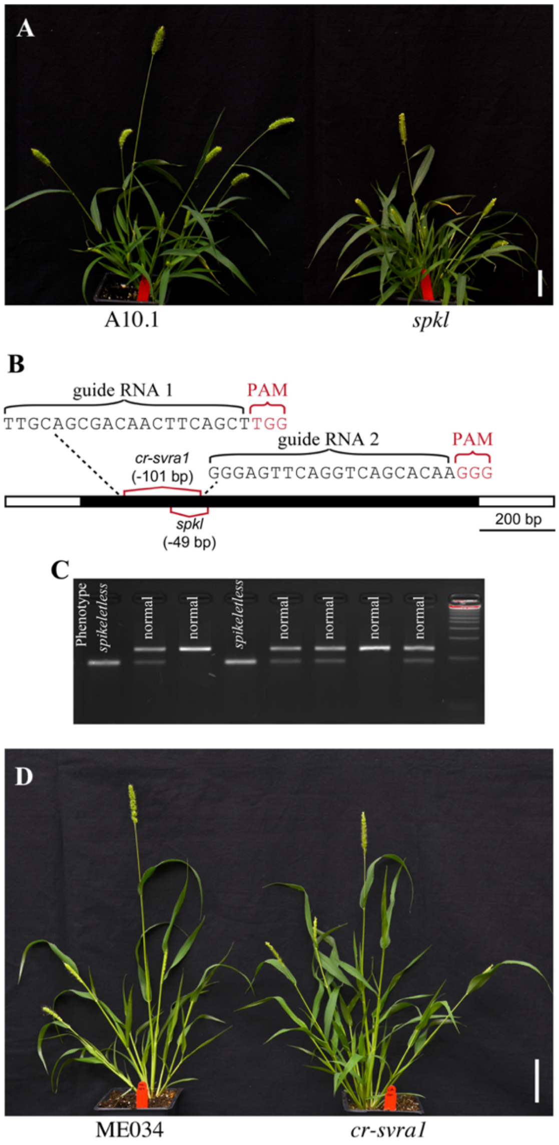


**Figure S7. Whole plant morphologies of *spkl* and edited *cr-svra1* and gene model of SvRa1 showing locations of deletions in *spkl* and *cr-svra1*.**

(A) Plant morphology of parental genotype, A10.1 (left), and *spkl* (right) mutant. (B) Schematic diagram of *Spkl/SvRa1* gene model showing location and sequences of two guides used to generate CRISPR (CR)-CAS9 based deletion. The 49 bp deletion in *spkl* and 101 bp deletion *cr-svra1* are also shown. PAM: Protospacer Adjacent Motif. (C) Genotyping results showing homozygous 49 bp deletion in *spkl* co-segregate with spikeletless phenotype. Plants containing at least one copy of the reference allele showed normal panicles. (D) Plants of *cr-svra1* (right) compared to parental genotype, Me034 (left).

**Scale bars:** 5 cm (A, D).

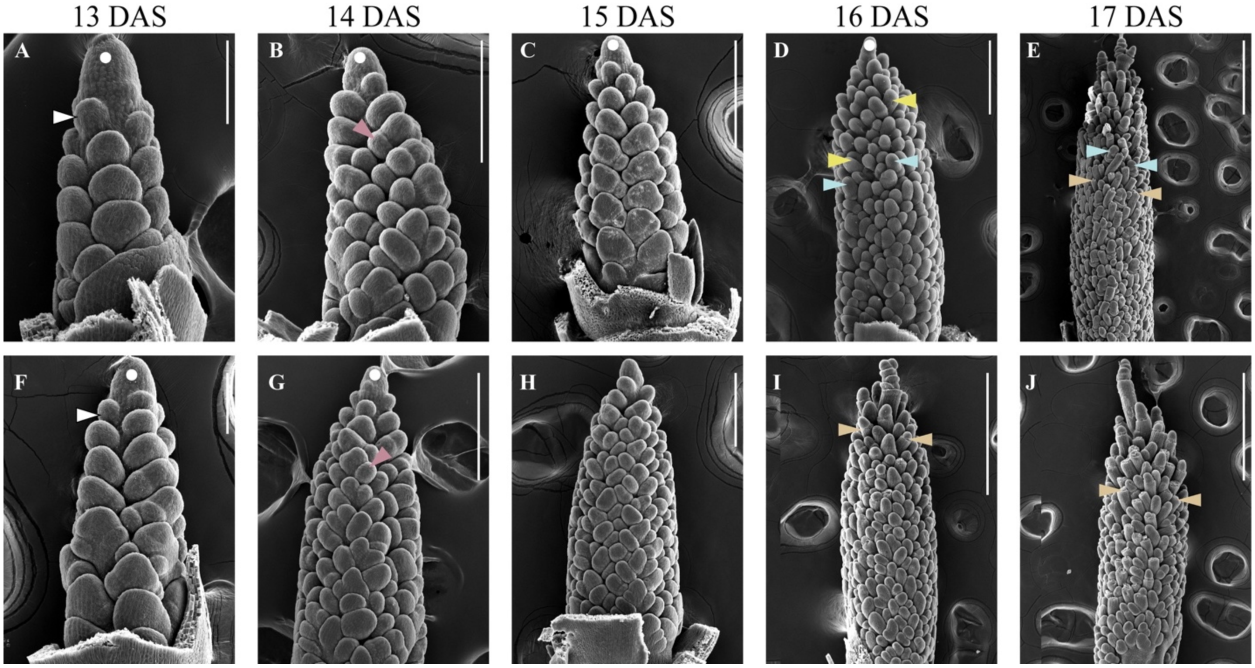


**Figure S8.** Scanning electron microscopy images showing developmental progression of A10.1 and *spkl* inflorescence primordia.

(A-C, F-H) In both A10.1 and *spkl*, primary branch meristems (BMs, white arrow) and higher-order BMs (pink arrow) formed around 13-15 days after sowing (DAS) and inflorescence meristem (IM, white dot) acquired determinate fate around 15 DAS.

(D) In A10.1 panicles, some BMs near the tip elongated (yellow arrow) and a few others appeared bulged (blue arrow).

(E) Around 17 DAS, most BMs in A10.1 differentiated to form SM (blue arrow) or bristles (gold arrow).

(I) In *spkl,* bristles with indented ring (gold arrow) around meristem were visible near panicle top by 16 DAS.

(J) At 17 DAS in *spkl,* only bristles were formed. Here, gold arrows indicate where meristem tip appears to break off in the mature bristle.

**Scale bars:** 100 µm (A, F), 200 µm (B, H), 250 µm (C-D, G), 500 µm (E, I-J).

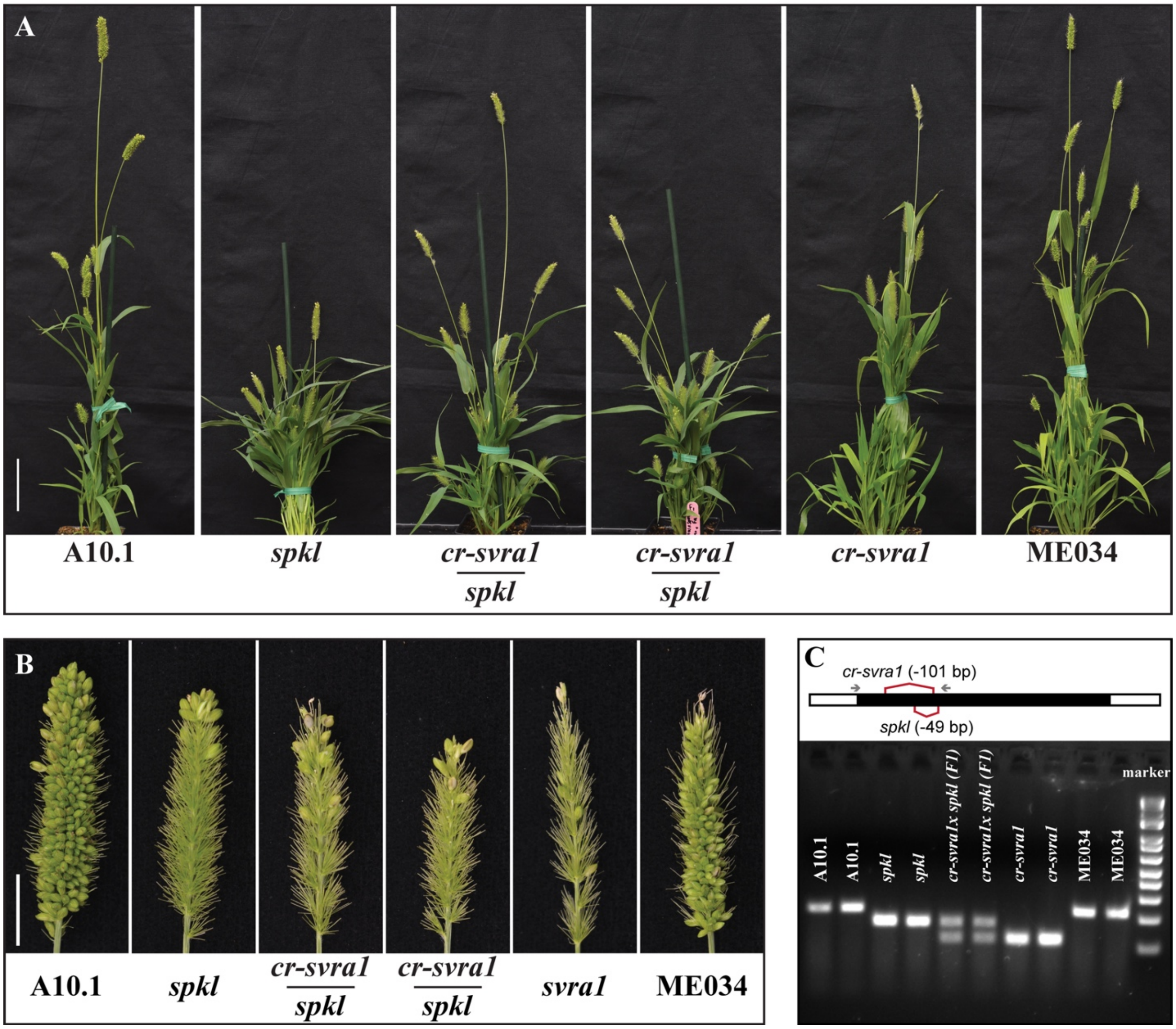


**Figure S9. Plants, panicles and genotyping of F1s obtained from genetic cross between *spkl* and *cr-svra1*.** (A) Plant morphology and (B) panicles and (C) genotyping of parental and F1 plants. In gene model (in C), deletions in *spkl* and *cr-svra1*, and location of primers (gray arrow) used in genotyping are shown.

**Scale bars:** 10 cm (A), 1 cm (B).

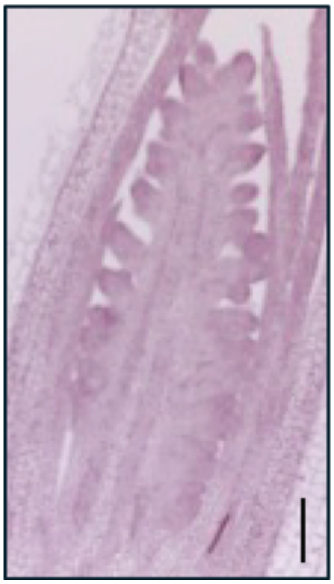

**Figure S10.** *In-situ* hybridization using *SvRa1* sense probe showed no detectable signal in panicle collected from A10.1 plants at 17 days after sowing. Scale: 100 µm.

Figure S11

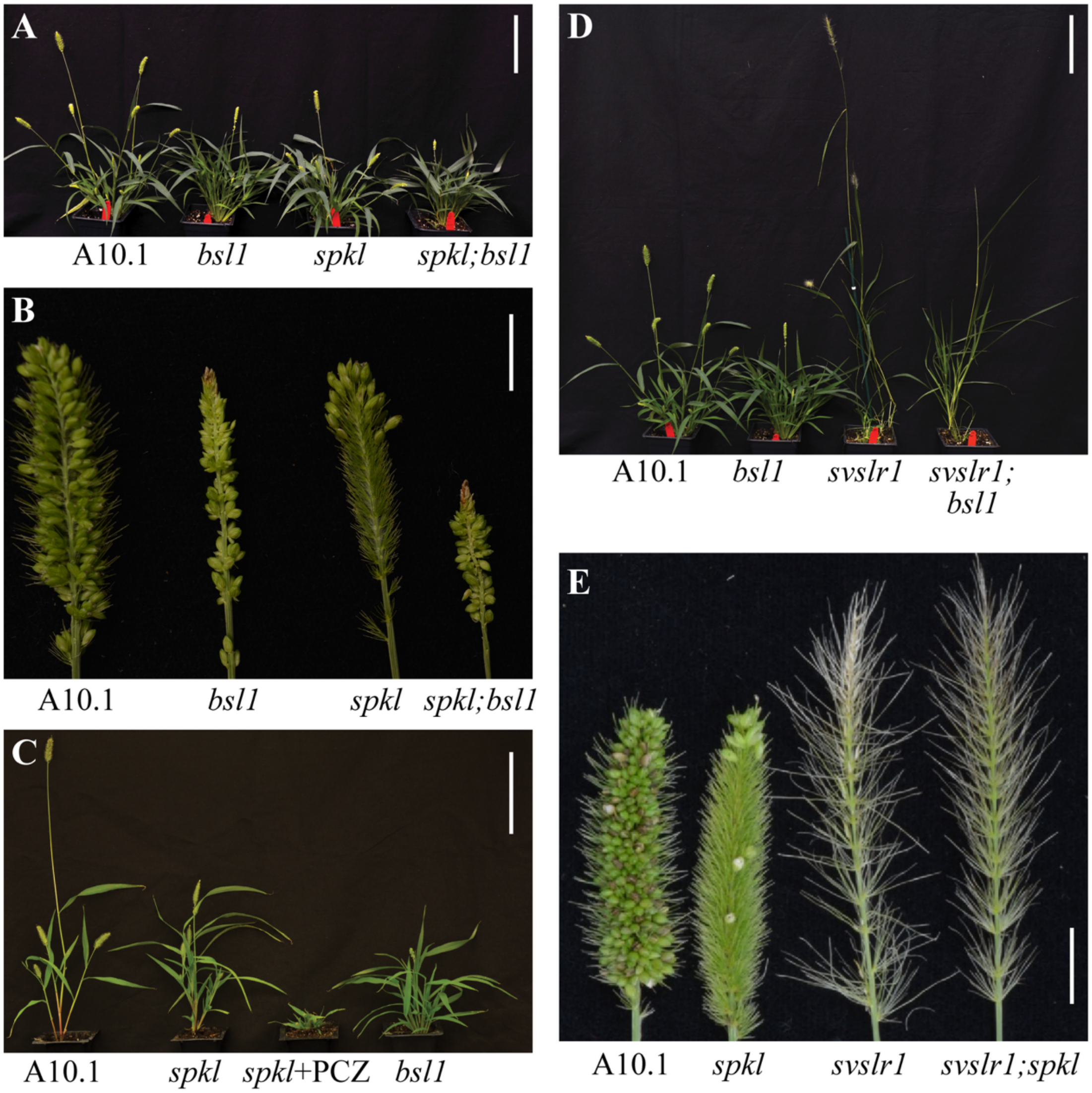


**Figure S11**. **Comparing whole plant and panicle morphologies of A10.1, single mutants (*bsl1, spkl* and *svslr1*) and double mutants (*spkl;bsl1, svslr1;bsl1, svslr1;spkl*); and effect of Propiconazole (PCZ) treatment on whole plant morphology of *spkl*. (A)** Plants and **(B)** panicle cross sections of A10.1*, bsl1* (*bsl1-1* allele from Yang et al. 2018)*, spkl and spkl; bsl1.* **(C)** Plant morphology of Propiconazole (PCZ)-treated *spkl* compared to A10.1*, bsl1* and *spkl.* **(D)** Plant phenotypes of A10.1, *bsl1*, *svslr* 1and *svslr1;bsl1.* **(E)** Panicles of A10.1*, spkl, svslr1* and *svslr1*;*spkl.*

**Scale bars:** 10 cm (A, C, D), 1 cm (B, E).

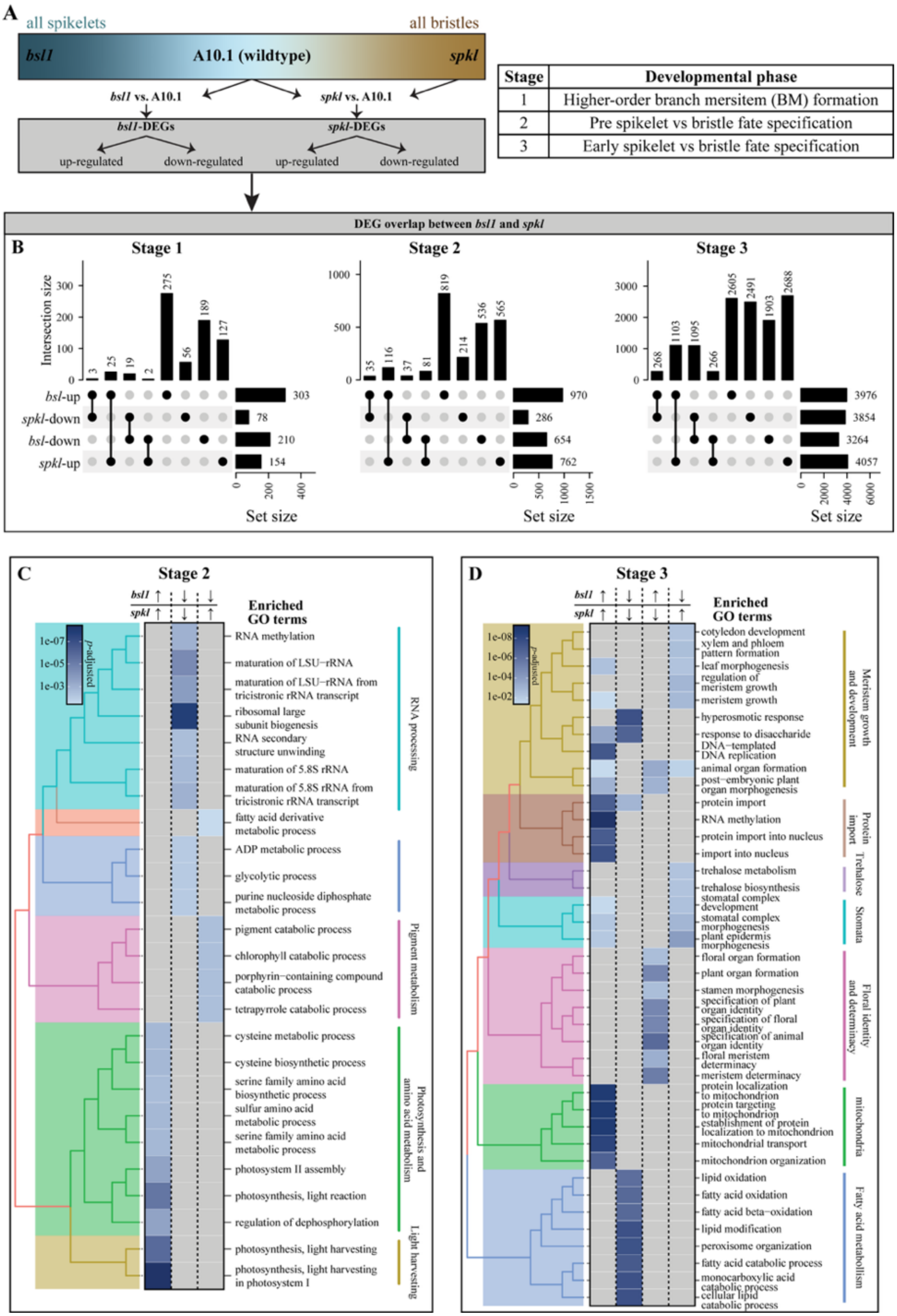
A
all spikelets
all bristles
bsl1
A10.1 (wildtype)
spkl
bsl1 vs. A10.1
spkl vs. A10.1
bsl1-DEGs
spkl-DEGs
up-regulated
down-regulated
up-regulated
down-regulated
Stage | Developmental phase
1 | Higher-order branch mersitem (BM) formation
2 | Pre spikelet vs bristle fate specification
3 | Early spikelet vs bristle fate specification
DEG overlap between bsl1 and spkl
B
Stage 1
Stage 2
Stage 3
Intersection size
3 25 19 2 275 56 189 127
35 116 37 81 819 214 536 565
268 1103 1095 266 2605 2491 1903 2688
bsl-up
spkl-down
bsl-down
spkl-up
303 78 210 154
970 286 654 762
3976 3854 3264 4057
Set size
C
Stage 2
bsl1
spkl
Enriched GO terms
p-adjusted
1e-07
1e-05
1e-03
RNA methylation
maturation of LSU−rRNA
maturation of LSU−rRNA from tricistronic rRNA transcript
ribosomal large subunit biogenesis
RNA secondary structure unwinding
maturation of 5.8S rRNA
maturation of 5.8S rRNA from tricistronic rRNA transcript
fatty acid derivative metabolic process
ADP metabolic process
glycolytic process
purine nucleoside diphosphate metabolic process
pigment catabolic process
chlorophyll catabolic process
porphyrin−containing compound catabolic process
tetrapyrrole catabolic process
cysteine metabolic process
cysteine biosynthetic process
serine family amino acid biosynthetic process
sulfur amino acid metabolic process
serine family amino acid metabolic process
photosystem II assembly
photosynthesis, light reaction
regulation of dephosphorylation
photosynthesis, light harvesting
photosynthesis, light harvesting in photosystem I
RNA processing
Pigment metabolism
Photosynthesis and amino acid metabolism
Light harvesting
D
Stage 3
bsl1
spkl
Enriched GO terms
p-adjusted
1e-08
1e-06
1e-04
1e-02
cotyledon development
xylem and phloem pattern formation
leaf morphogenesis
regulation of meristem growth
meristem growth
hyperosmotic response
response to disaccharide
DNA−templated DNA replication
animal organ formation
post−embryonic plant organ morphogenesis
protein import
RNA methylation
protein import into nucleus
import into nucleus
trehalose metabolism
trehalose biosynthesis
stomatal complex development
stomatal complex morphogenesis
plant epidermis morphogenesis
floral organ formation
plant organ formation
stamen morphogenesis
specification of plant organ identity
specification of floral organ identity
specification of animal organ identity
floral meristem determinacy
meristem determinacy
protein localization to mitochondrion
protein targeting to mitochondrion
establishment of protein localization to mitochondrion
mitochondrial transport
mitochondrion organization
lipid oxidation
fatty acid oxidation
fatty acid beta−oxidation
lipid modification
peroxisome organization
fatty acid catabolic process
monocarboxylic acid catabolic process
cellular lipid catabolic process
Meristem growth and development
Protein import
Trehalose
Stomata
Floral identity and determinacy
mitochondria
Fatty acid metabolism

**Figure S12. Schematic representation of RNA-seq analysis comparing transcriptomes of A10.1, *spkl* and *bsl1* inflorescence primordia and GO term enrichment analysis of DE gene sets.** (A) Schematic representation of experimental setup. Inflorescence primordia at three developmental stages were collected from A10.1, *spkl* and *bsl1* plants. Mutants *spkl* and *bsl1* were compared to reference genotype A10.1 at corresponding stages to identify differentially expressed (DE) genes: up- and down-regulated. (B) Upsets plots showing overlap of DE genes between *spkl* and *bsl1*. (C-D) Tree plots of Gene Ontology (GO) terms enriched among genes that are DE in both mutants at (C) stage 2 and (D) stage 3. The arrows represent up- (↑) and down- (↓) regulated genes. Tree plots of top GO terms were generated using pairwise_termsim function in clusterprofiler package in R.

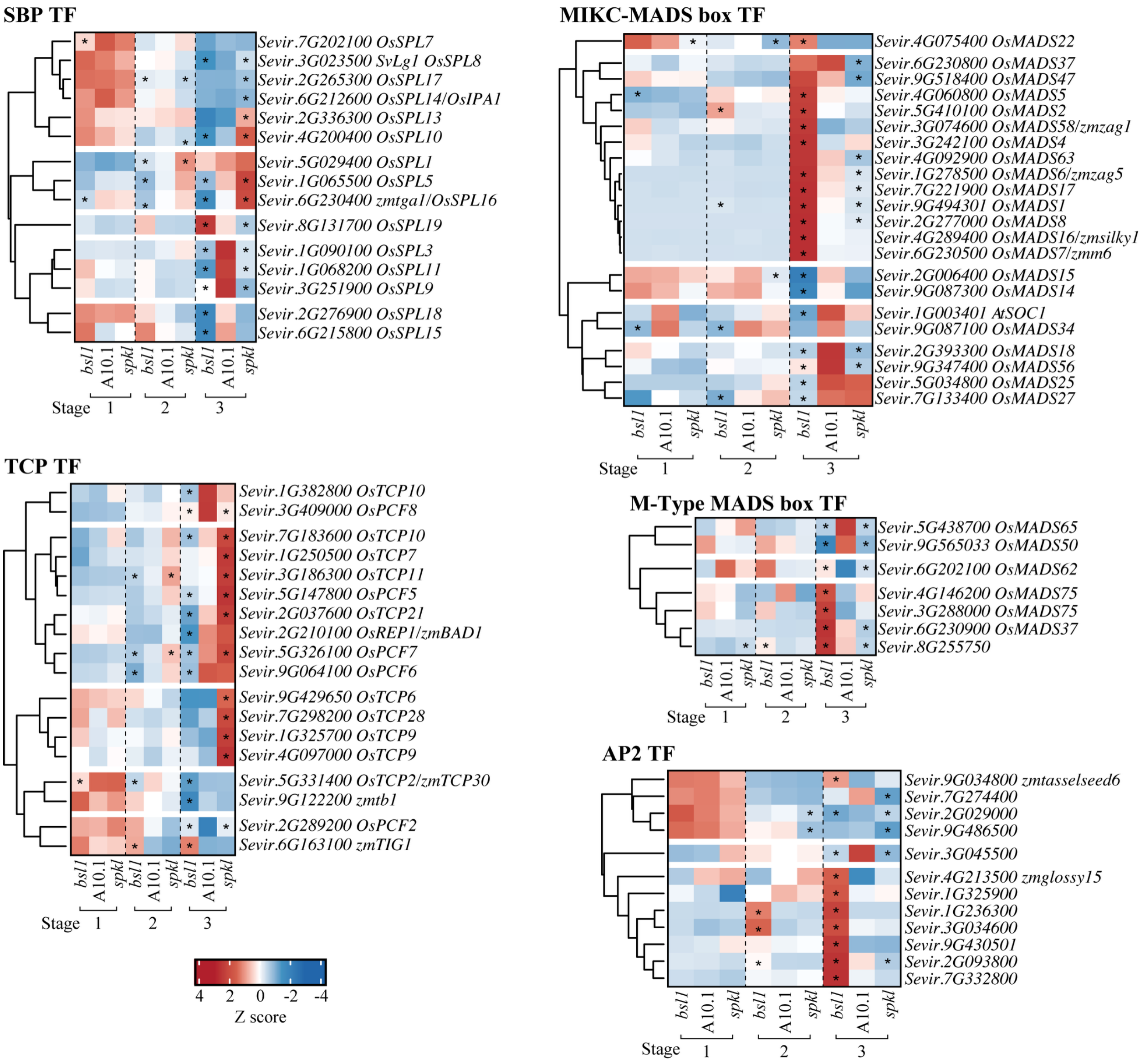


**Figure S13. Heatmaps showing expression profiles of selected transcription factor (TF) families in *bsl1*, A10.1, and *spkl* inflorescence primordia.** Here, z-scores represent scaled TPM values. Gene names for rice (*Os*), maize (*zm*), and Arabidopsis (*At*) orthologs are noted. Here, * indicate significant differential expression compared to A10.1 at same stage.

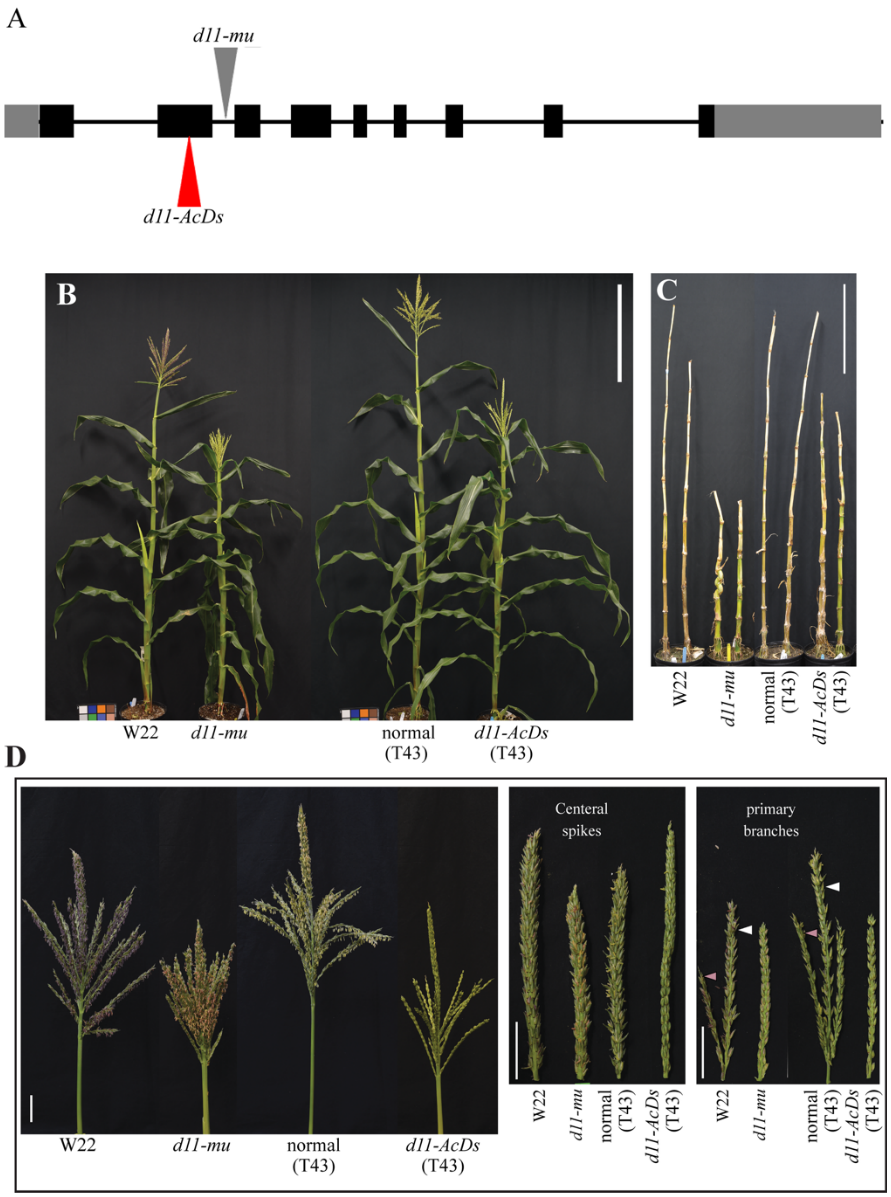


**Figure S14. Whole plant and tassel phenotypes of maize *dwarf11* (*d11*), *Bsl1* ortholog, alleles: *d11-mu* and *d11-AcDs*.** (A) Maize *d11* gene model showing insertion sites in *d11-mu* and *d11-AcDs* alleles. Here, *d11-mu* and *d11-AcDs* alleles are in W22 and T43 backgrounds, respectively. (B) Plants, (C) nodes and internodes, (D) tassels with peduncle (left), unbranched central spikes (middle) and 1$^{st}$ tassel primary branches (right) from W22, *d11-mu* (W22 background); and segregating normal (T43) and *d11-AcDs* (T43) plants. Here, white arrow: primary branches; pink arrow: secondary branches.

**Scale bars:** 50 cm (B-C), 5 cm (D).

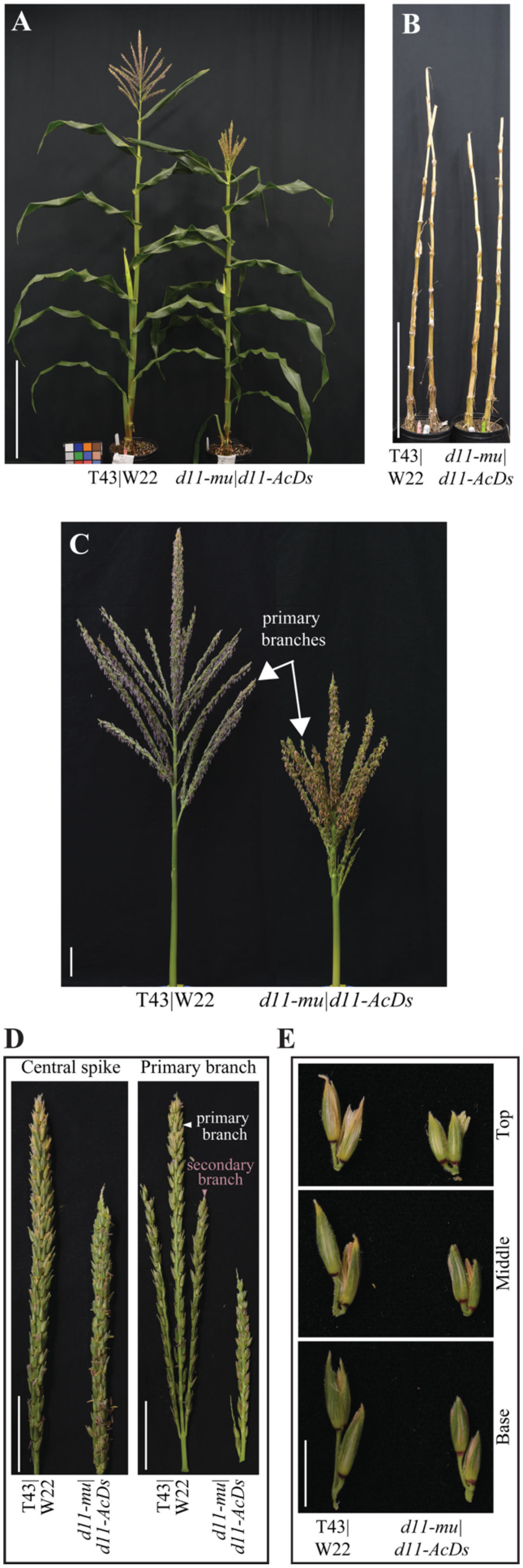


**Figure S15. Phenotypic characterization of F1s obtained from genetic cross between two alleles of *d11*: *d11-mu* and *d11-AcDs*.** (A-B) F1 between *d11-mu* and *d11-AcDs* showed (A) semi-dwarf plants and (B) shortened internodes compared to parental F1. (C) Tassels from Mutant-F1 (*d11-mu* X *d11-AcDs*) were smaller, and (D) showed shorter central spikes and reduced secondary branching (black arrow). (E) Spikelets and pedicels in mutant-F1 were shorter than parental F1. Scale: 50 cm (A-B), 5 cm (C-D), 1 cm (E).

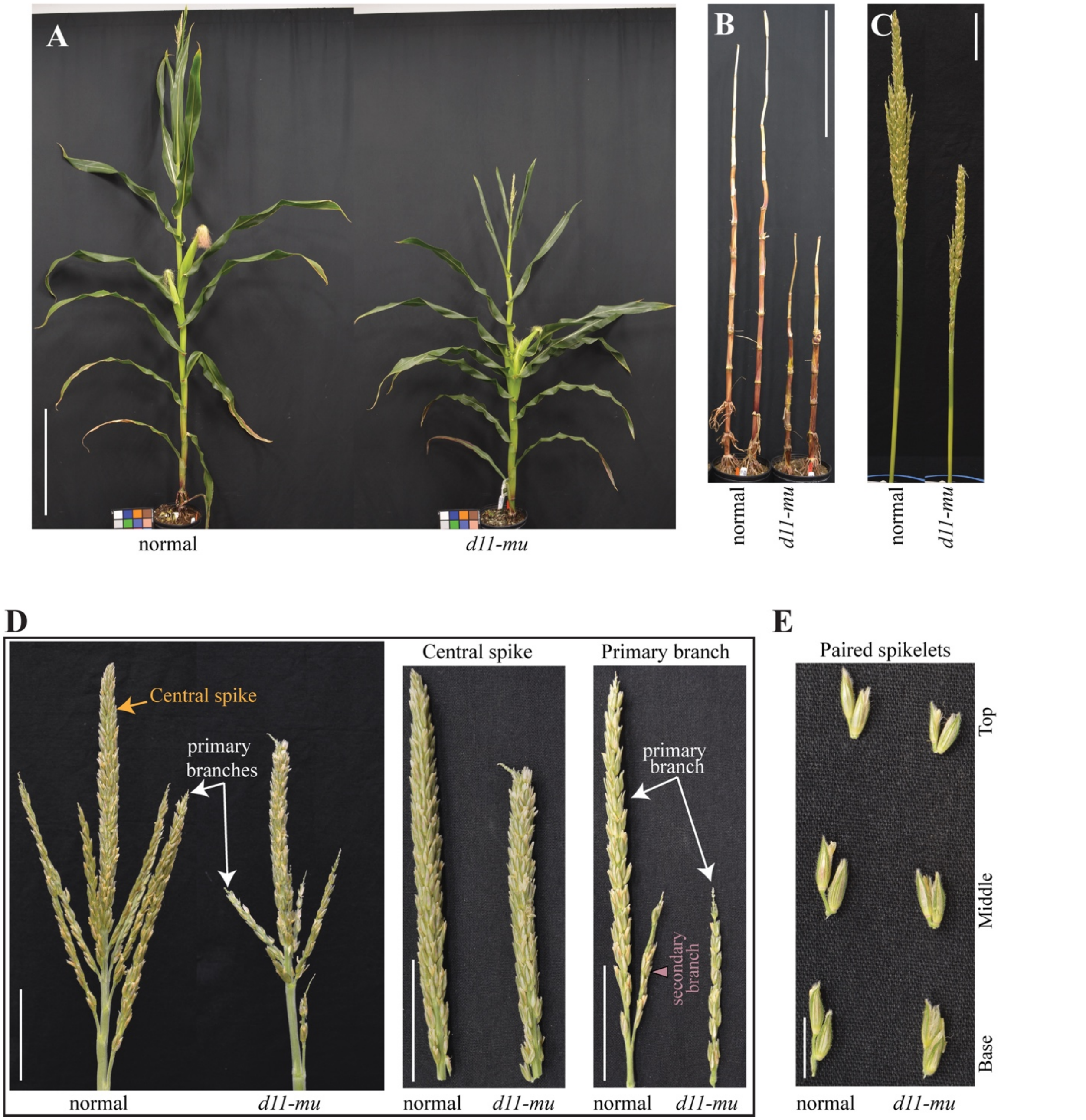


**Figure S16.** The *d11-mu* (W22) was backcrossed to B73 four times to introgress *d11- mu* into B73 background. Compared to normal plants, *d11-mu* showed (A) semidwarf plant (B) shortened internodes and (C-D) produced smaller tassels with shorter central spikes. (D) Tassels from *d11-mu* showed reduction in (white arrow) primary and secondary (pink arrow) branching (E) Compared to normal plants, *d11-mu* showed reduction in pedicel and spikelet length.

**Scale bars:** 50 cm (A-B), 5 cm (C-D), 1 cm (E).

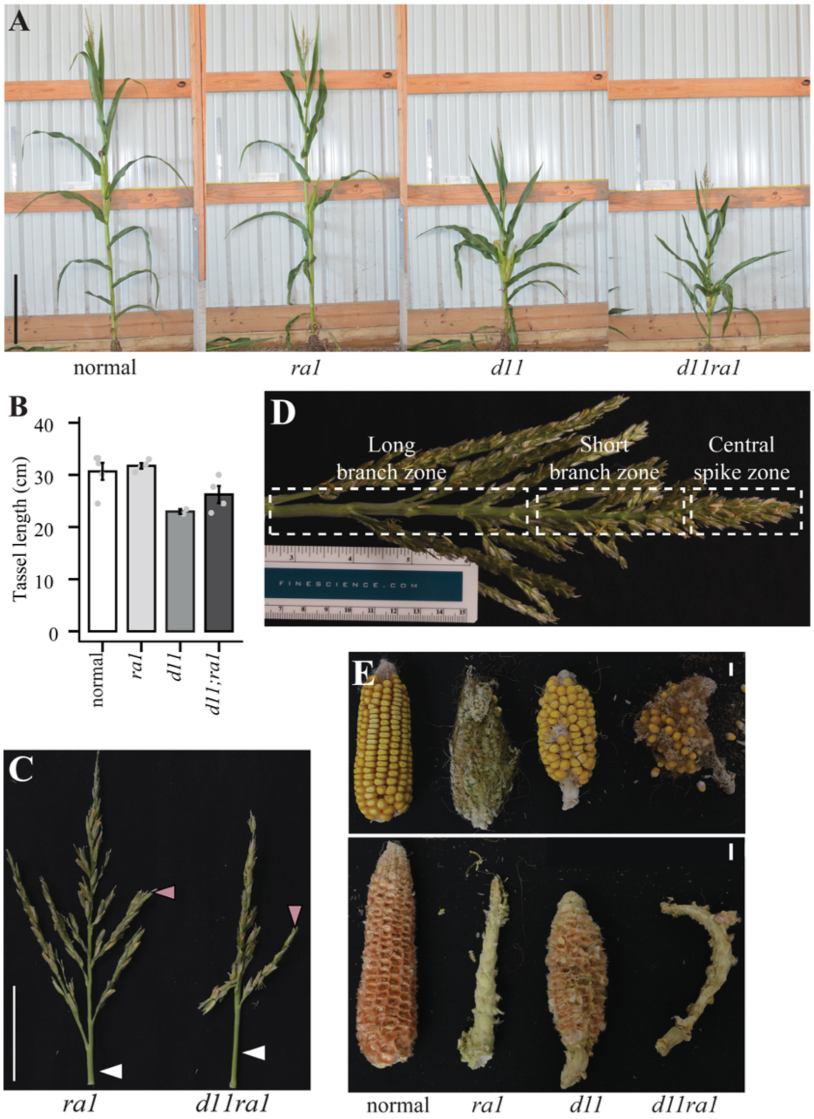


**Figure S17. Plant, tassel and ear phenotypes of normal, *d11-mu, ra1-R and d11-mu;ra1-R* plants obtained from genetic cross between *d11-mu* (B73 background) and the *ra1* reference allele (*ra1-R*; B73 background).** (A) Both the *d11* single mutant and *d11;ra1* double mutant showed semi-dwarf stature. (B) Both *d11* and *d11;ra1* showed reduction in tassel length compared to non-mutant (normal) and *ra1* individuals in field-grown plants (year 1). (C) Primary branches (white arrows) of *d11;ra1* produced fewer secondary branches (pink arrows) than *ra1*. (D) Distinct zones of *ra1* tassels: long branch, short branch, and central spike. (E) Ears from segregating plants show branching in *ra1* and *d11;ra1* (top). Naked cobs after removal of kernels (in normal and *d11*) and branches (in *ra1* and *d11;ra1*) are shown below.

**Scale bars:** 50 cm (A), 5 cm (C), 1 cm (E).

**Table S1. Measurements of panicle traits for A10.1, *spkl, svslr1,* ME034 and cr-*svra1.*** Two-tailed t-test was used to compare *spkl* and *svslr1* to corresponding parental genotype, A10.1; and *cr-svra1* to parental genotype, ME034. Here, * $p < 0.05$, ** $p < 0.01$, *** $p < 0.001$, **** $p < 0.0001$, ns: non-significant.

| | n | Panicle length (cm) | No. of clusters[1] per cm of panicle ($cm^{-1}$) | No. of spikelets per cluster[1] | No. of bristles per cluster[1] | Total no. of structures[2] per cluster[1] |
|---|---|---|---|---|---|---|
| A10.1 | 28 | 3.42 ± 0.07 | 15.52 ± 0.54 | 7.29 ± 0.28 | 18.57 ± 0.62 | 25.86 ± 0.86 |
| *spkl* | 28 | 2.81 ± 0.06**** | 24.69 ± 0.92**** | 0.04 ± 0.03**** | 25.84 ± 1.13**** | 25.88 ± 1.14[ns] |
| *svslr1* | 20 | 4.31 ± 0.20*** | 8.62 ± 0.75**** | 0.14 ± 0.07**** | 10.54 ± 0.63**** | 10.68 ± 0.62**** |
| ME034 | 20 | 3.91 ± 0.14 | 11.1 ± 0.44 | 11.40 ± 0.53 | 15.84 ± 0.74 | 26.98 ± 1.25 |
| *cr-svra1* | 20 | 4.66 ± 0.13*** | 10.95 ± 0.55[ns] | 0.24 ± 0.06**** | 30.79 ± 0.92**** | 31.03 ± 0.9* |

[1] A cluster comprises all higher order branches originating from a primary branch meristem. Here, five clusters from equivalent sections of each panicle were used to collect average bristle and spikelet number per cluster of a panicle.

[2] Here, aggregate of spikelet and bristle number per cluster denotes total number of terminal structures per cluster.

**Table S2. Plant height and tiller count data for A10.1, *spkl, svslr1, bsl1, svslr1;bsl1* and *spkl;bsl1* collected at the flowering stage.** Multiple non-parametric t-tests with "bonferroni" correction for multiple testing were used to compare all genotypes. Different letters indicate means are significantly different from each other (*padj* < 0.01).

| | n | Plant height (cm) | Tiller count |
|---|---|---|---|
| A10.1 | 42 | $40.27 \pm 0.66^{c}$ | $17.05 \pm 0.63^{d}$ |
| *spkl* | 44 | $16.64 \pm 0.59^{e}$ | $33.75 \pm 1.11^{b}$ |
| *svslr1* | 32 | $66.10 \pm 2.42^{a}$ | $3.97 \pm 0.32^{f}$ |
| *bsl1* | 44 | $23.08 \pm 0.64^{d}$ | $27.77 \pm 0.66^{c}$ |
| *svslr1; bsl1* | 27 | $51.96 \pm 1.90^{b}$ | $8.70 \pm 0.54^{e}$ |
| *spkl; bsl1* | 23 | $15.33 \pm 0.79^{e}$ | $48.7 \pm 1.72^{a}$ |

**Table S3. Plant height and tiller count data for parental genotype ME034 and *cr-svra1***. Here, two-tailed t-test was used to compare *cr-svra1* to ME034. **** $p <=$ 0.0001, ns: non-significant.

| | **n** | **Plant height (cm)** | **Tiller count** |
|---|---|---|---|
| ME034 | 28 | 50.97 $\pm$ 1.65 | 13.82 $\pm$ 0.45 |
| *cr-svra1* | 28 | 47.6 $\pm$ 1.51$^{ns}$ | 27.61 $\pm$ 0.94**** |

**Table S4.** RNA-seq based transcript abundance (Transcripts Per Million, TPM) for all *Setaria viridis* (*Sv*, V2.1) genes in A10.1, *bsl1*, and *spkl* inflorescence primordia across the three developmental stages. (A) Mean of biological replicates and (B) values for all replicates are provided. *Sv* annotation file (Sviridis_500_v2.1.annotation_info.txt" ) containing functional annotations and orthologs from rice and Arabidopsis was obtained from phytozome. Maize orthologs were extracted from phytozome biomart using *Sv* (V2.1) gene identifiers as query. The classical maize gene annotations are from MaizeGDB. Transcription factor (TF) annotations are from plantTFDB (https://planttfdb.gao-lab.org/).

**Table S5.** Differentially expressed (DE, *p*-adj < 0.05) genes in *bsl1* and *spkl* compared to parental genotype, A10.1. The annotations were obtained from *Setaria viridis* genome annotation file (V2.1) in phytozome.

**Table S6.** Differentially expressed (DE, *p*-adj < 0.05) genes with contrasting expression changes ($log_2$-fold change relative to A10.1) in *bsl1* and *spkl* at (A) stage 2 and (B) stage 3.

**Table S7.** Gene Ontology terms enriched among gene sets that are DE in both *bsl1* and *spkl* mutants. The four differentially expressed (DE) genes sets are: (1) up-regulated in both *bsl1* and *spkl*, (2) down-regulated in *bsl1* and *spkl*, (3) up-regulated in *bsl1* and down-regulated in *spkl,* (4) down-regulated in *bsl1* and up-regulated in *spkl.* GO term enrichment analysis for (A) stage 2 and (B) stage 3 is presented.

**Table S8.** Enrichment of (A) transcription factors (TFs) and (B) TF families among genes sets that are DE in both mutants at stage 3. These DE gene sets are: (1) up-regulated in both *bsl1* and *spkl*, (2) down-regulated in *bsl1* and *spkl*, (3) up-regulated in *bsl1* and down-regulated in *spkl,* (4) down-regulated in *bsl1* and up-regulated in *spkl.* Following code in R was used to test the enrichment: p_value <- phyper(k-1, K, N-K, n, lower.tail = FALSE; Fold_enrichment <- (k/n)/(K/N).

**Table S9.** (A) Plant and (B) tassel traits for *d11* mutants and corresponding non-mutants. Here, *d11-mu* allele is in W22 and B73 background, and *d11-AcDs* is in T43 background. Two-tailed t-test was used to compare *d11-mu* (W22) to W22, *d11-AcDs* (T43) to normal (T43), *d11-mu* (B73) to normal (B73) and mutant-F1 (*d11-mu-W22/d11-AcDs-T43*) to T43/W22.

**Table S10** Phenotypic analysis of *ra1, d11, d11;ra1* and normal plants obtained from cross between *ra1* and *d11-mu* (B73). (A) Two-way ANOVA was used to test the interaction between factors *d11* and *ra1.* (B) Tukey HSD test was used to test differences between means.

**Table S11:** Primers used in the study.
